# Impaired value estimation over space explains turn-induced freezing of gait in Parkinson’s disease: A cortico-basal ganglia-inspired reinforcement learning model

**DOI:** 10.64898/2026.10.01.755546

**Authors:** Neermita Bhattacharya, Sandeep Sathyanandan Nair, V. Srinivasa Chakravarthy, Vignesh Muralidharan

## Abstract

Parkinson’s disease (PD) gait is characterized by reduced walking speed, short shuffling steps, increased double stance time, and increased step time variability. The most debilitating symptom in PD is freezing of gait (FOG), a context-dependent locomotion arrest, mostly occurring while encountering turns, obstacles, and narrow pathways. FOG is more pronounced during turning, a complex task that challenges motor planning and execution. Here, we simulate this phenomenon using a computational model inspired by the cortico-basal ganglia circuitry, based on reinforcement learning (RL) principles. In the model, the agent was trained to learn a value profile while navigating a path that involved turning. To simulate PD conditions, we clamped the temporal difference (TD) error, typically associated with dopaminergic signaling. Our model results show that compared to healthy controls (HC), PD non-freezers and PD freezers exhibit a reduction in turning velocity and step length, alongside an attenuated compensatory step-width widening at the sharpest turn, more pronounced in PD freezers, who additionally show increased step time variability. These kinematic impairments worsen during tighter turns. Ultimately, this model implicates FOG as a problem of impaired estimation of value over space and could potentially represent a generalized deficit in gait impairments in PD.

## INTRODUCTION

Freezing of gait (FOG) is a very common symptom associated with Parkinson’s disease (PD) characterized by a sudden inability to move forward, which is particularly experienced while initiating a movement, encountering a turn, or facing an obstacle such as a narrow pathway ahead^1^. Being one of the most disabling symptoms, FOG is also one of the most poorly understood. FOG significantly impairs mobility, increases the risk of falls, and negatively impacts the overall quality of life for PD patients^2^. Understanding the causes and finding effective intervention strategies for FOG is crucial in improving the management and treatment of PD. Turning is one of the most prominent causes that induces freezing in PD patients^3,4^. On monitoring gait in a natural setting, the percentage of freeze episodes caused by turning is higher compared to other triggering events even under medicated conditions^5^.

Turning requires a shift of motor programs, i.e., from normal gait where the swing amplitude of both legs are similar, to altering the amplitude of one leg to achieve a turn towards the right or left. This would require proper set-shifting, a critical factor of executive control which is often lacking in patients who experience freezing (PD-FOG)^6^. Negotiating a turn requires the ability to dynamically shift the body’s center of mass (CoM) from its existent base of support while moving in a new direction^7^. Therefore, a range of compensatory mechanisms is employed by healthy individuals while encountering a turn. They include increased step widths, slower turning speeds, larger turning angles, and a craniocaudal sequence of turning, which help maintain balance^7^. Analogous to the classic speed-accuracy trade-off in motor control, gait involves a speed-balance trade-off critical for preventing postural instability during turns^8,9^. However, PD patients experience problems in controlling gait timing and fail to increase step width to the same extent as healthy controls, especially while negotiating sharper turns leading to freezing episodes^10^. Hence the turning strategies adopted and their effectiveness in maintaining stability is the differentiating factor between healthy controls and PD.

Healthy controls (HC) adapt to the complexity of turns by following a craniocaudal sequence (head, thorax, then pelvis). In contrast, PD-FOG patients rotate all segments simultaneously^11^. Another characteristic of PD-FOG patients negotiating a turn is shorter step lengths they take to complete them^12^. HC, on the contrary, takes bigger strides with the leg that is moving towards the direction of the turn. A study conducted by Chee et al.^13^ shows that irrespective of change in directions, it is typical of PD-FOG to experience gradual shortening of step lengths as time progresses. It remains unclear whether this shortening of step length while simultaneously encountering a turn results in balance issues causing the freeze. Reduced speed and increased turning duration are other features associated with turning in PD-FOG compared to PD non-FOG^14^. In addition to the step length, PD-FOG adopts a step out and mixed strategy for turning while HC adopts a crossover strategy. This strategy adopted by PD-FOG can be associated with the failure to increase step width^10^. Moreover, the turning angles play a significant role in the extent of the freeze with sharper turns aggravating the situation with PD patients showcasing more step time variability and increased freeze times during turns^10^.

In literature, there are various mechanisms and contributing factors involved in turn freezing in PD. Some of the significant factors include i) the motor complexity involved^6,15^, ii) postural instability^7^, iii) cognitive load^16,17^, and iv) sensorimotor integration^6^. Turning involves a complex coordination of muscle groups and joints, requiring more precise motor control than straight walking. PD patients, due to impaired motor control and bradykinesia, find it challenging to execute these movements smoothly. The additional balance demands can trigger FOG as the patient attempts to compensate^15^. Turning increases cognitive load because it requires spatial navigation and planning. Cognitive deficits in PD, including issues with executive function and attention, exacerbate the difficulty of turning, leading to FOG. Effective turning requires the integration of sensory feedback with motor actions. PD disrupts this integration, causing delays and inaccuracies in the motor response, which can result in FOG. Turning has also been coupled to a secondary task like color identification to study the effects of dual tasking^16^. Evidence from these studies warrants understanding of turning in PD-FOG and the influence of turning environment on gait.

However, a generalized framework explaining FOG in different contexts is still elusive. Computational models estimating the value function over space have shown promise for understanding the neural mechanisms of FOG^17–19^. In our earlier work, a reinforcement learning (RL) model of the cortico-basal ganglia circuitry demonstrated that FOG can emerge from degraded spatial value representations, replicating the velocity dips and step length reductions seen when PD-FOG navigate narrow doorways^18^. This spatial evaluation framework was later expanded to incorporate utility-based decision-making, effectively capturing how high cognitive load in processing complex Stroop cues while walking can increase expected uncertainty and trigger motor arrests^17^. Since spatial and cognitive complexities distort the internal value of space, many aspects of turn freezing can also be captured using a similar approach. In this study, we extend our previously proposed RL-based cortico-basal ganglia model to explain turn freezing in PD. Specifically, we use it to interpret the findings from two clinical studies: the first examined the influence of turning sharpness on HC, PD non-FOG, and PD-FOG^10^; the second observed PD-FOG and PD non-FOG performing a virtual reality gait task where they exhibited motor arrests during turn execution^20^. Our model successfully predicts key spatial and temporal gait changes, including turning velocity, step length, step width, and step time coefficient of variation and attributes it to impaired estimation of value over space.

## MATERIALS AND METHODS

### The Turning Environment

The proposed model simulates a subject’s navigation through a continuous path containing a defined curve, computing the resultant kinematic profiles — including turning velocity, step length, step width, and step time CV, along the track (Fig. 1a). During this task, the agent encounters turns of varying sharpness (e.g., 0°, 60°, 90°, 120°, and 180°) and must successfully navigate the curve to reach the end of the corridor without colliding with the lateral boundaries (Fig. 1b). To ensure that the total distance walked by the agent remains consistent across these different turning scenarios, the length of the straight track segments is adjusted based on the turn angle (ϕ_*track*_) as described in Table 1. The input parameter ϕ _*track*_ defines the angular deviation in radians of the initial and final track segments from the central horizontal axis. The total actual turn angle executed by the agent is 2ϕ _*track*_. To mirror clinical observations, the agent was evaluated on input parameters of 0°, 30°, 45°, 60°, and 89° (converted to radians), which correspond to the turning severities of 0°, 60°, 90°, 120°, and 178° (180°) respectively. The 89° parameter was specifically used to approximate the 180° turn. This prevents track boundary overlaps in the generation of the simulation’s geometric environment. The simulated 178° condition is referred to as the 180° turn hereafter for simplicity.

**Table 1.**
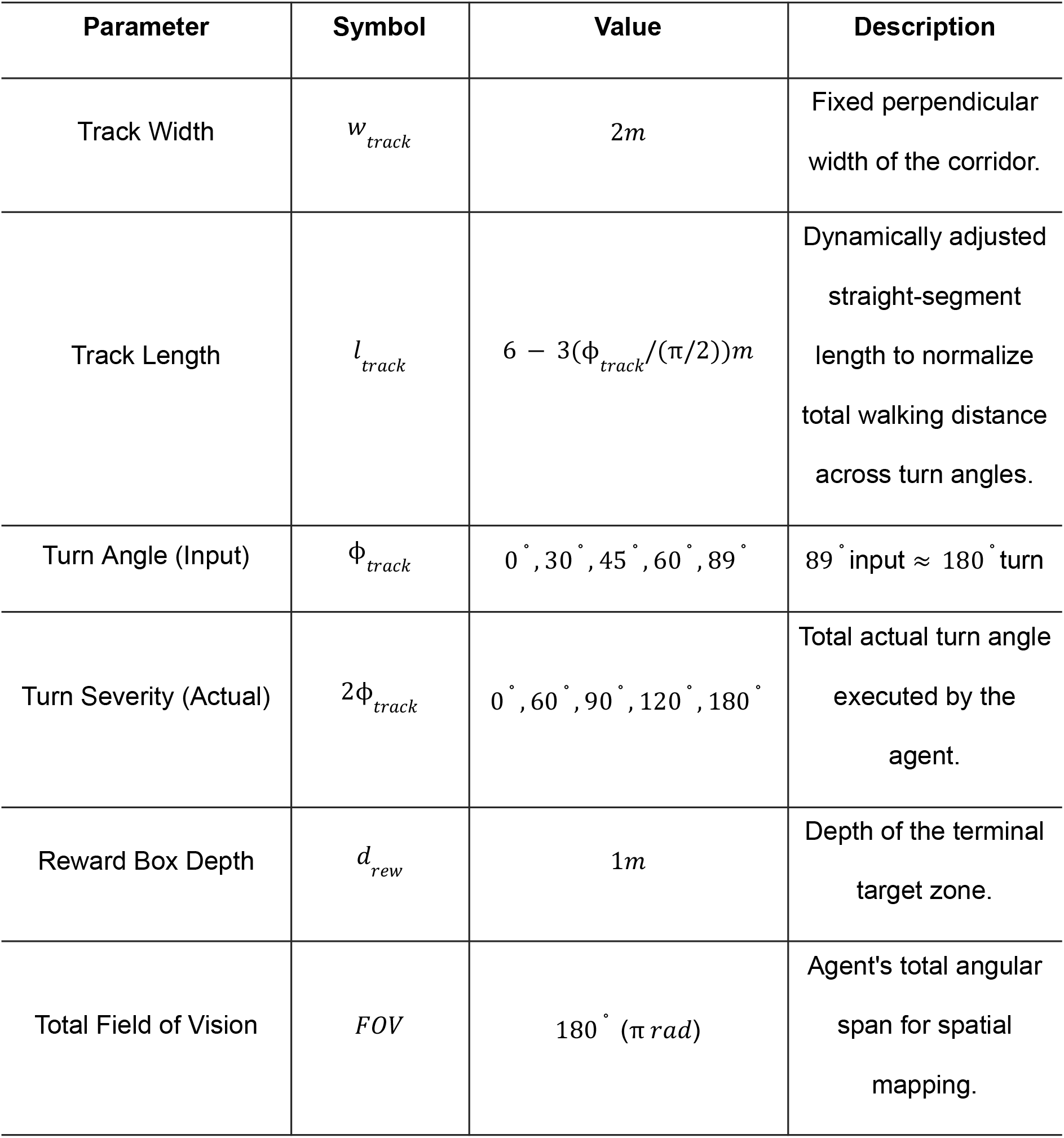
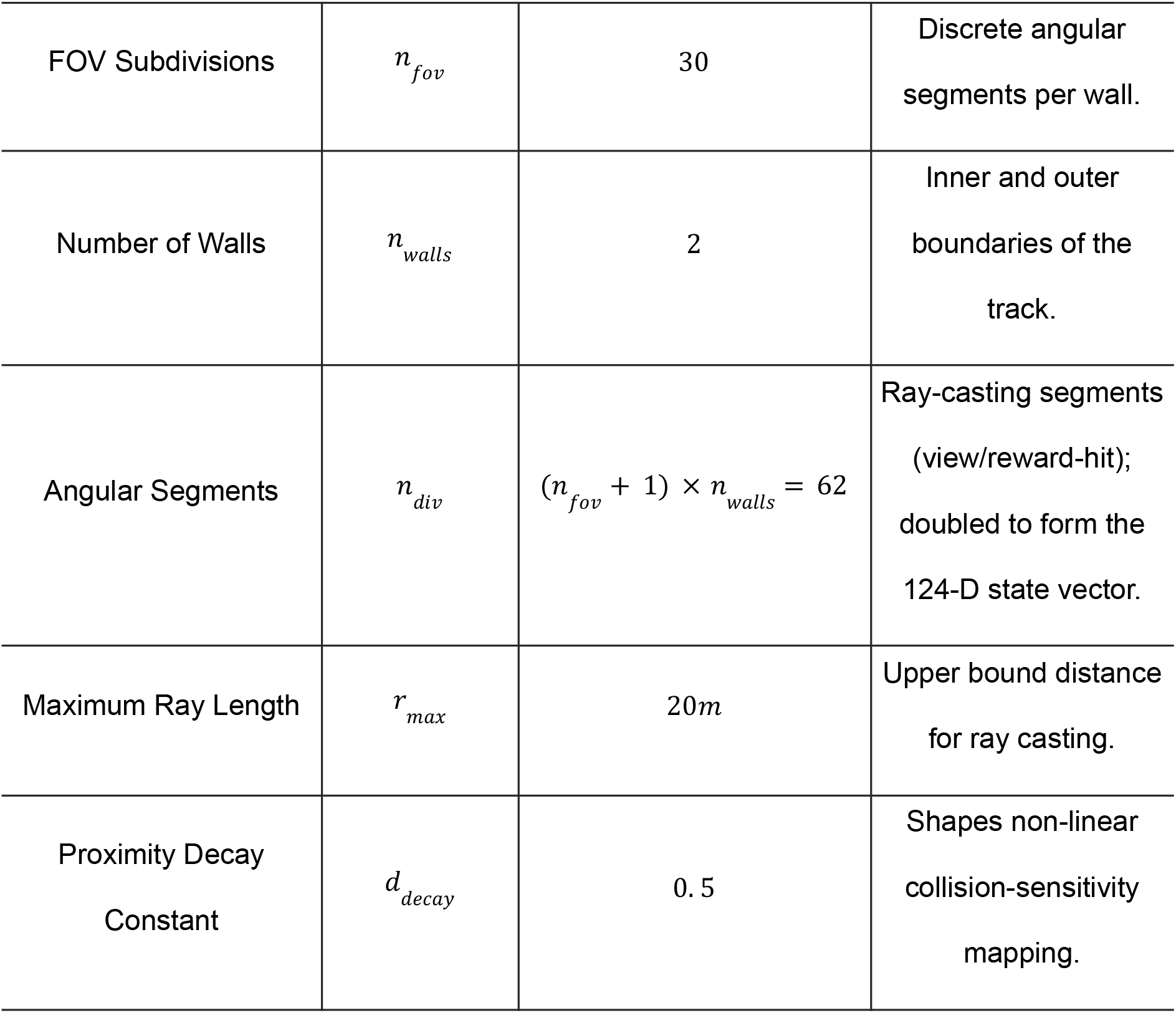
**Environment Parameters** — geometry and sensory configuration of the turning track and the agent’s visual field.

**Figure 1.**
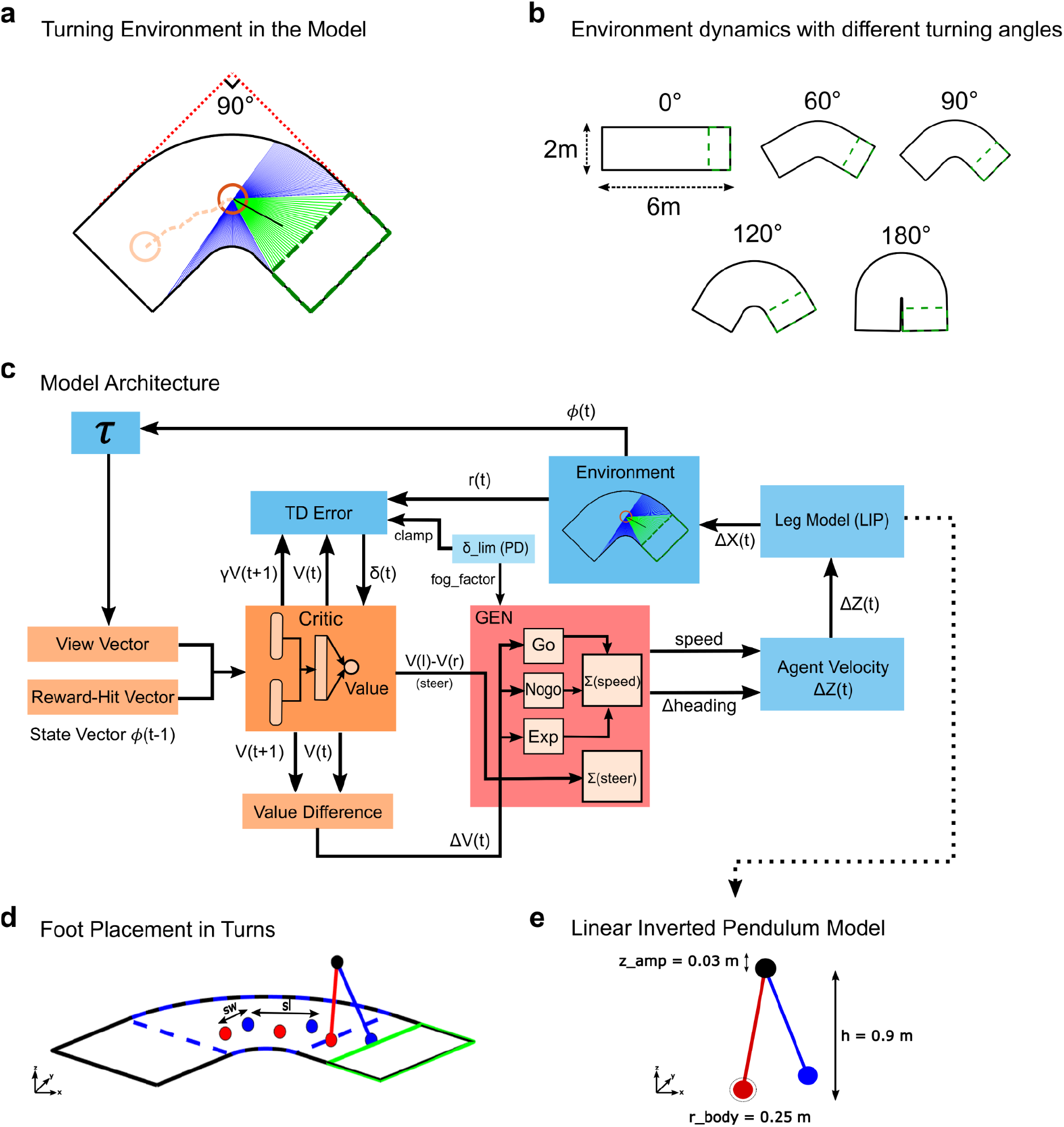
**Overview of the Turning Environment and Model Architecture:** (a) Schematic of the simulated environment showing an agent approaching a 90° turn; the agent navigates a corridor toward a target reward zone (green dashed box) while negotiating the turn. The green lines indicate the rays in the agent’s FOV that reach the reward box; blue lines indicate the rays that hit the walls. (b) Simulated environment at different turn angles. (c) Model architecture schematic showing information flow from the state vector through the Critic (value estimation), the GEN action-selection block (Go/Explore/No-go gains and value-gradient steering), the LIP leg model (gait generation), and back through the environment to close the reinforcement learning loop. (d) 3D foot placement along a 90° turn, showing step length (*sl*) and step width (*sW*). (e) 3D schematic of the Linear Inverted Pendulum (LIP) model, showing biomechanical parameters including center-of-mass height (*h*), vertical oscillation amplitude (*z*_*amp*_), and body radius (*r*_*body*_) defining the physical footprint for boundary-collision detection.

Completing a turn successfully is defined by reaching a designated target zone at the end of the track (Fig. 1a, green dashed box), which yields a positive reward. The agent’s movement speed is governed by an inherent speed-accuracy tradeoff^8,9,21,22^. High velocities, particularly during sharp turns, increase the required turning radius and the probability of boundary collisions. Consequently, through RL mechanisms, the agent learns an optimal policy to decelerate upon entering a turn, maximizing its reward by avoiding collision penalties. This computational framework is specifically utilized to capture and explain the empirical spatial and temporal gait modifications exhibited by PD subjects as they negotiate turns of different severities, directly mirroring clinical observations^10,16,23,24^.

### Computational Model

The proposed model architecture (Fig. 1c) consists of three main components: the cortico-basal ganglia (CBG) neural system, the kinematic agent module (Action Selection and Gait Generation), and the turning environment. All scripts were custom written in MATLAB R2026a (MathWorks, Natick, MA, USA) for the model. The cortical area compiles spatial information into a 124-dimensional state vector (ϕ(*t*)) and forwards it to the Critic block within the CBG model to compute the current state value *V*(*t*). The resultant Value Difference Δ*V*(*t*), which is the temporal change in expected reward, is sent to the Action Selection block. Action selection regulates the agent’s movement via two distinct pathways: a scalar speed controller driven by direct Go/No-go and Exploratory gains, and a value-gradient steering controller that probes directional headings to figure out how much to turn. The outputs of these pathways are integrated to update the agent’s velocity vector. Based on this continuous velocity and position data, the gait module estimates discrete kinematic metrics including step length, step width, turn velocity and step time (Supplementary Fig. S1). Simultaneously, upon acting on the environment the agent receives a reward *r*(*t*). This reward, combined with the Critic’s state evaluations, generates a temporal difference (TD) error δ(*t*) that trains the Critic network’s parameters over successive trials.

The full simulation pipeline was executed using 15 distinct random seeds (*rng*(1) − *rng*(15)), each generating 10 independent episodes. The resulting data from these 150 total episodes were subsequently combined for all kinematic, spatial, and behavioral analyses.

#### The State Vector Block

This module computes the agent’s spatial awareness state, allowing the agent to localize itself optimally within the track. Ray casting is utilized to detect the boundaries of the environment (the inner and outer walls) as well as the target reward zone. Each of the *n* discrete angular segments is assigned a ray angle θ *i*, calculated as an offset from the agent’s current heading direction, evenly distributed across the agent’s total field of view (Eq.1) For each *i*-th angular segment directed at angle θ *i* over a distance *r*, the continuous ray coordinates 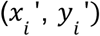 are mapped using standard polar-to-Cartesian transformations (Eq. 2). These coordinate pairs make the view ray vector for each segment at a given time *t*. The specific parameters defining the agent’s field of vision, ray limits, and angular resolution are detailed in Table 1.

The state of the agent is represented by the concatenation of two distinct properties extracted from 62 discrete visual rays mapping the environment: a view vector (proximity vector) and a reward-hit vector. The 62-element proximity vector is estimated by applying an exponential decay function to the distance of each ray intersecting the track boundaries (Eq. 3), making the agent highly sensitive to imminent collisions. Conversely, the 62-element reward-hit vector evaluates target localization; rays successfully intersecting the reward box are coded as 1s, and the rest as 0s. These two 62-element arrays are concatenated to form the final 124-dimensional state vector input ϕ(*t*) sent to the Critic block.

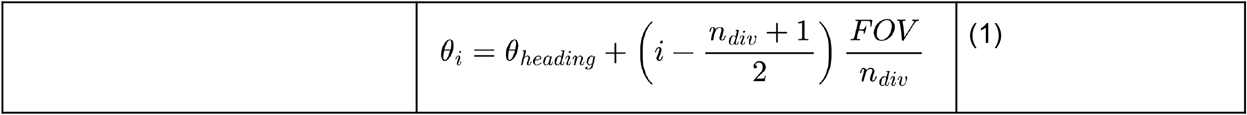

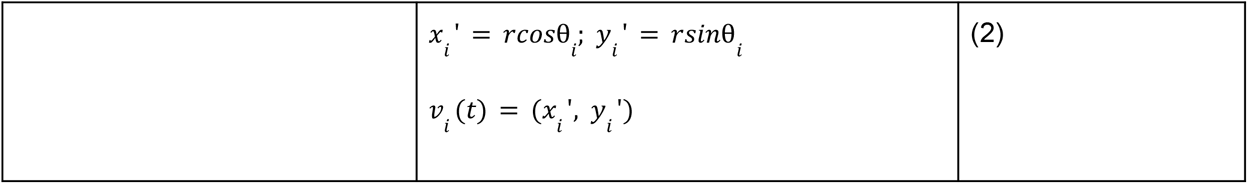

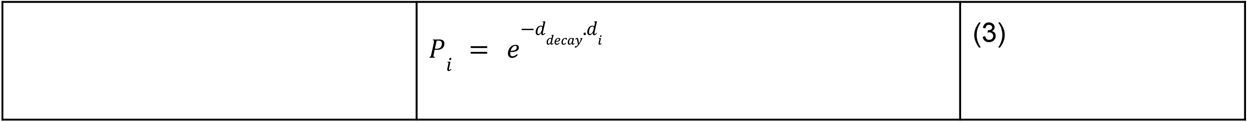

where *P* _*i*_ is the proximity vector, and *d* _*i*_ is the distance from the agent to a boundary along ray *i*.

#### The Critic Block

The Critic block in the model mimics the cortico-basal ganglia dynamics of computing and learning the value function *V*(*t*), an approach taken in several previous studies^17,18,25,26^. In our case, we modeled the Critic block as a feed-forward multi-layer perceptron (MLP) receiving the state vector ϕ(*t*) as input. To capture the complexity of the state space, the network utilizes two hidden layers (consisting of 32 and 64 neurons, respectively, Table 2) with a hyperbolic tangent (*tanh*) activation function, and a single linear output node that estimates the scalar value function. The Critic block also receives input from the temporal difference block, which utilizes the current state value *V*(*t*), the discounted value of the subsequent state *V*(*t* + 1), and the reward from the environment (*r*(*t*)) to estimate the temporal difference (TD) error δ(t). This TD-error acts as the dopaminergic teaching signal and is propagated backward to train the parameters of the Critic network via gradient descent^26^. Finally, the value information is utilized to compute Δ*V*(*t*) — representing the step-by-step change in expected future rewards, which functions as the primary driving signal for the Action Selection block.

**Table 2.** **Critic Parameters** — architecture and learning parameters of the value-estimating neural network.

| Parameter | Symbol | Value | Description |
| --- | --- | --- | --- |
| Hidden Layer<br>Sizes | — | [32, 64] | Units in the two hidden layers of the MLP. |
| Output<br>Dimension | — | 1 | Single linear output node estimating $V(t)$. |
| Activation<br>Function | $f(x)$ | $\tanh(x)$ | Non-linearity used in hidden layers. |
| Discount Factor | $\gamma$ | 0.9 | Temporal reward discounting, Eq. (7). |
| Initial Learning<br>Rate | $\eta_0$ | 0.01 | Initial learning rate, Eq. (6). |
| Learning Rate<br>Decay Constant | — | 500 | Decay time constant ($n_{trials}$) |
| Learning Rate | $\eta$ | $\eta_0 \cdot e^{(-trial/n_{trials})}$ | Decaying Learning Rate |
| Condition Flag | $pdcond$ | 0 (HC) / 1 (PD) | Switches whether the TD-error clamp and downstream GEN scaling<br>are applied. |
| TD Clamp (PD<br>Non-FOG) | $\delta_{lim}$ | 0.008 | Upper TD clamp, Eq. (27). |
| TD Clamp (PD<br>FOG) | $\delta_{lim}$ | 0.001 | Upper TD clamp, Eq. (27). |
| Medication<br>Offset | $\delta_{med}$ | 0 | Additive TD-error offset<br>(zero at baseline). |
| Update Rule | — | Mini-batch<br>gradient descent | Weight update method<br>during the first 450 trials. |

The forward pass of the Critic network computing the value function *V*(*t*) is defined in Eq. 4.

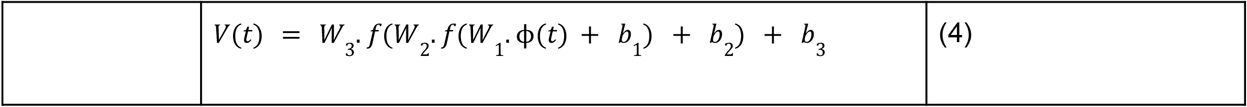

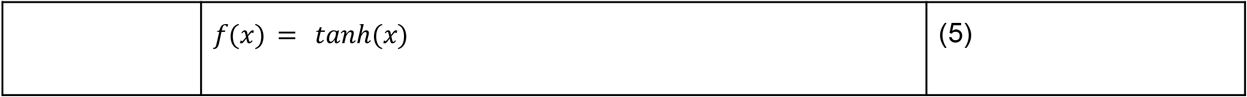

Where *W n* and *b n* represent the weight matrices and bias vectors for each layer, and the non-linear activation function *f*(*x*) is the hyperbolic tangent (Eq. 5). The network parameters are updated using backpropagation driven by the learning rate η and the TD-error δ(*t*):

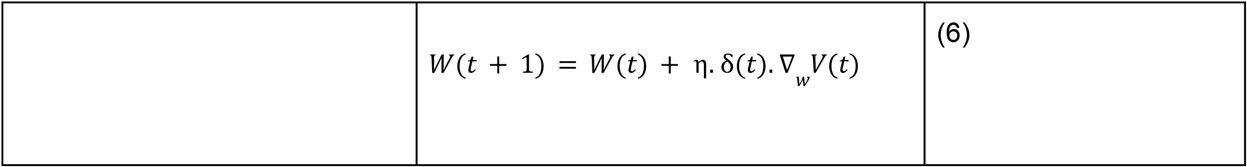

The temporal difference (TD) error is formulated utilizing the discount factor (γ):

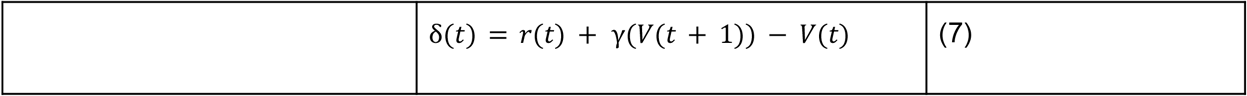

To ensure stable convergence, the Critic network parameters are updated using mini-batch gradient descent during the training phase (the first 450 trials), utilizing the accumulated TD-errors from the entire trial. Also, η in Eq. 6 is not static; it is decayed over successive trials using an exponential schedule denoted in Table 2. This allows the network to transition smoothly from rapid initial exploration to fine-tuned exploitation as the agent learns the track.

#### The Action Selection/GEN block

Action selection in the CBG circuitry is achieved through a competitive process between the direct and indirect pathways^27,28^. The Action Selection block works on the Go/Explore/No-go (GEN) policy, where exploration plays a key role in learning optimal movement trajectories. While traditional models combine these signals into a single vector, this implementation separates the movement into a scalar speed controller and a value-gradient steering mechanism. This allows the agent to dynamically decelerate and steer smoothly through sharp turns. The primary driving signal for the GEN policy is Δ*V*(*t*), defined in Eq. 8.

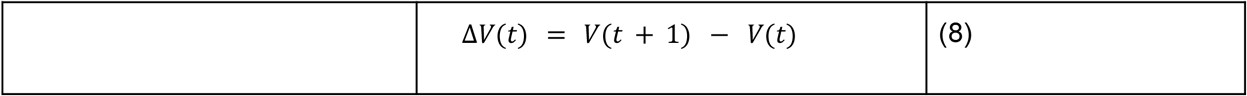

The direct pathway (Go) and indirect pathway (No-go) gains are computed using sigmoid activation functions driven by Δ*V*(*t*). The exploration term *E* enables stochastic hill-climbing over the value function when the gradient is weak, decaying naturally as the agent learns (controlled by variance σ^2^).

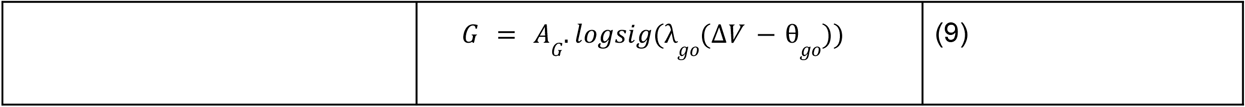

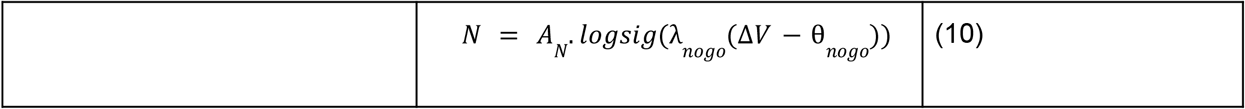

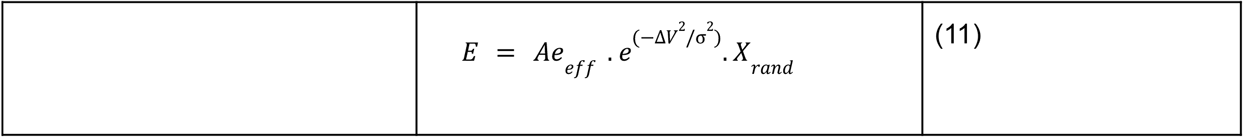

where *A* _*G*_, *A* _*N*_, and *Ae* _*eff*_ are the pathway amplitudes (Table 3), θ represents the activation thresholds, and *X* _*rand*_ is a 2D uniform random noise vector. The full vector magnitude ||*E*|| is applied to the forward speed target, while its first scalar component *E*_*steer*_, is applied to the steering signal. The exploration amplitude *Ae*_*eff*_ and the exploration decay variance σ are not static; sharper turns begin with a larger initial *Ae*_*eff*_ to ensure higher exploration (Eq. 12), while σ decays faster (Eq. 13) to stabilize movement. During the testing phase (*trial* > 450),this variance-based exploration is disabled and replaced with a minor fixed amplitude uniform noise ± 0. 05 to evaluate the learned exploitation policy.

**Table 3.** **GEN (Action Selection) Parameters** — gains and thresholds governing the Go/Explore/No-go action-selection mechanism, including their scaling under simulated turn sharpness and dopaminergic depletion.

| Parameter | Symbol | Value | Eq. Ref |
| --- | --- | --- | --- |
| Go Pathway Amplitude | $A_G$ | 3 | (9) |
| No-go Pathway Amplitude | $A_N$ | 3 | (10) |
| Go Slope Parameter | $\lambda_{go}$ | 1 | (9) |
| No-go Slope Parameter | $\lambda_{nogo}$ | − 100 | (10) |
| Go Activation Threshold | $\theta_{go}$ | 0 | (9) |
| No-go Activation Threshold<br>(HC) | $\theta_{nogo}$ | 0.0 | (10) |
| Exploration Amplitude<br>Base/Slope | — | 1, 2 | (12) |
| Exploration Variance<br>Floor/Scale/Decay | — | 0.1, 0.5, 500 | (13) |
| Testing-Phase Exploration<br>Noise | — | $\pm 0.05$ | — |
| Steering Probe Base/Slope | — | 0.15, 0.55 | (17) |
| Steering Gain Base/Slope | — | 0.3, 0.6 | (23) |
| Turning Rate Base/Slope | — | 2.0, 5.0 | (24) |
| Maximum Steering Signal<br>Base/Slope | — | 1.0, 0.5 | (20) |
| Post-Turn Steering Damping | — | $\times 0.30$ | — |
| Sharp-Turn Speed Scaling<br>Slope | — | $- 0.01$ | (14) |
| Reference TD Clamp | $\delta_{lim\_ref}$ | 0.01 | (28) |
| PD Go Scaling Coefficient | — | 0.25 | (30) |
| PD No-go Scaling Coefficient | — | 0.25 | (31) |
| PD No-go Threshold Shift<br>Coefficient | — | 0.15 | (32) |

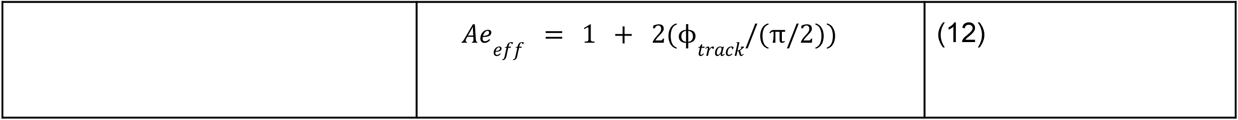

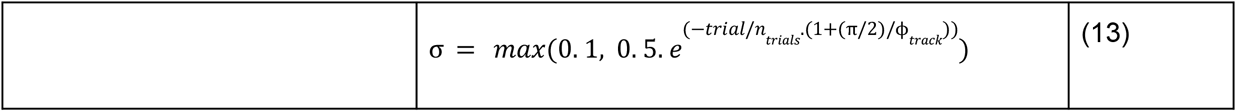

These pathway signals converge to form the agent’s target action speed. To ensure fluid, biomechanically realistic movement, the actual agent speed |*v*| is smoothly updated towards the target speed using a first-order leaky integrator governed by a rate constant λ_*v*_ and time step *dt*. For sharp maneuvers (≥90°), an additional speed-scaling penalty (Eq. 14) is applied at each step to the final velocity vector, reducing speed proportionately to the turn severity to simulate biomechanical caution inside the turn:

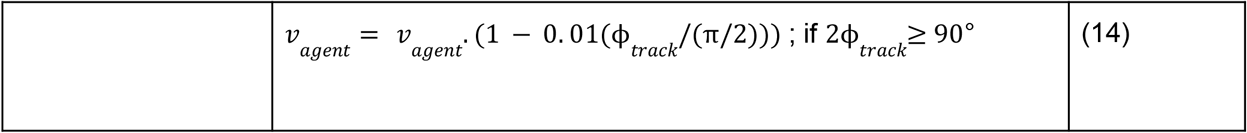

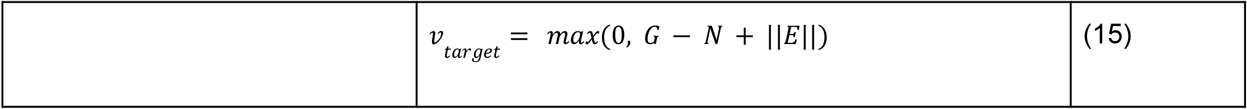

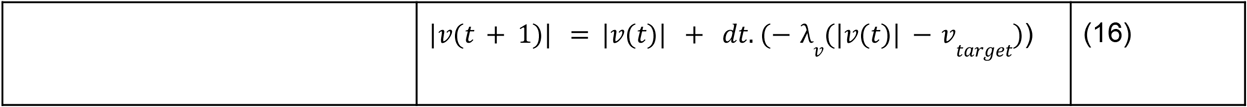

where *dt* is an integration parameter (Table 4).

**Table 4.** **Model Hyperparameters** — general simulation and training settings

| Parameter | Symbol | Value | Description |
| --- | --- | --- | --- |
| Total Episodes | $N_{epi}$ | 150 ($10 \times 15seeds$) | Total independent episodes pooled for analysis. |
| Random Seeds | — | $rng1 - 15$ | Seeds for the episode sets. |
| Trials per Episode | $n_{trials}$ | 500 | Trials simulated per episode. |
| Maximum Steps per Trial | $n_{steps}$ | 400 | Max agent steps per trial before termination. |
| Training/Testing Split | $train\_threshold$ | 450 | Trial index separating training/testing. |
| Integration Time Step | $dt$ | $0.02s$ | Update step for speed and steering. |
| Rate Constant | $\lambda_v$ | 2.5 | Rate constant of leaky-integrator speed controller, Eq. (16). |

Simultaneously, the agent determines its heading through a value-gradient steering controller. Instead of moving blindly, the agent probes the Critic network for the expected value of turning slightly to the left (*V*_*left*_) or right (*V*_*right*_). The angular deviation (δ_*steer*_) used for these directional probes adaptively widens in proportion to the track angle, allowing the agent to look further into sharper corners.

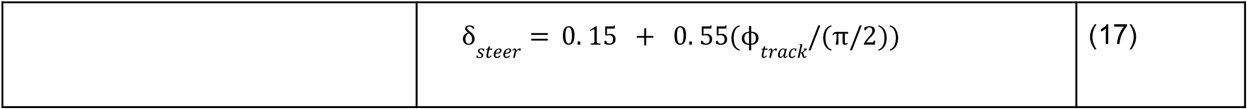

The difference between these spatial values creates a steering gradient *S*:

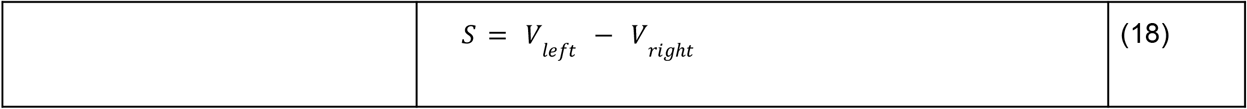

This gradient, combined with the exploratory signal, continuously updates a steering action signal *u*_*s*_.

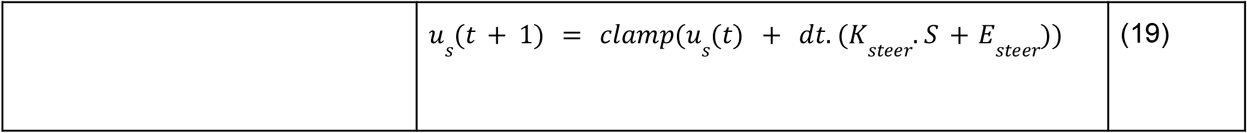

To prevent erratic oscillations, this steering signal is clamped to a maximum limit that scales with the track angle shown in Eq. 20. Furthermore, after successfully exiting the turn, the steering signal is dampened by 70% to stabilize a straight-line trajectory.

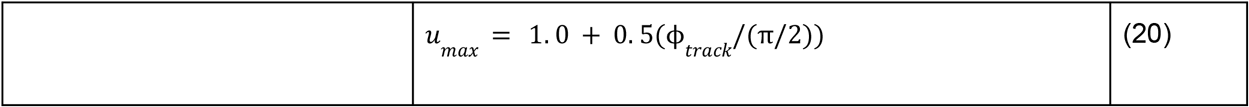

where − *u*_*max*_ ≤ *u*_*s*_ ≤ *u*_*max*_.

The agent’s physical heading vector is then updated by applying a rotation matrix, where the angle of rotation (Δθ) is proportional to the steering signal.

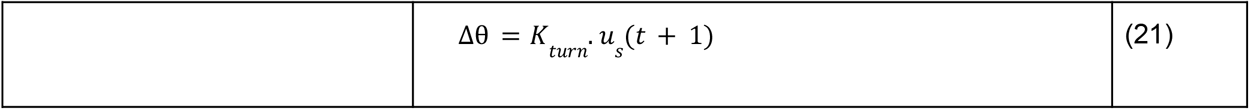

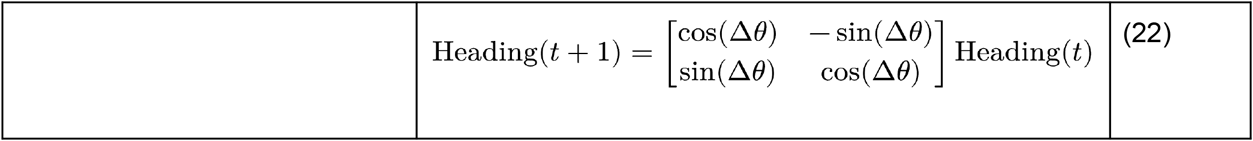

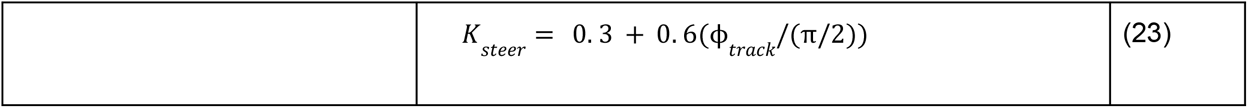

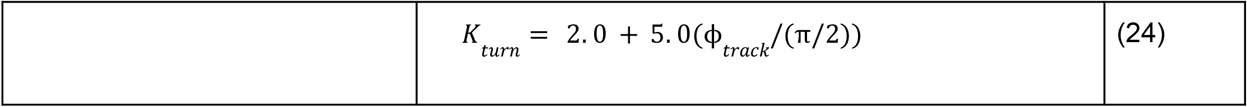

where the steering gain *K*_*steer*_ and turning rate *K*_*turn*_ dynamically scale with ϕ _*track*_ to allow sharper maneuvering in tighter corners.

Finally, the updated continuous velocity vector (*v*_*x*_, *v*_*y*_)is passed to a Linear InvertedPendulum (LIP) biomechanical model^29,30^ to generate discrete stepping metrics. Thetemporal timing of the steps (step time, *t*_*step*_) adapts inversely to the agent’s speed based on the natural frequency of the LIP model:

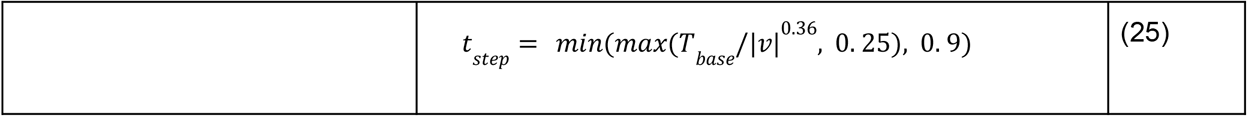

where *T*_*base*_ is the base half-period of the pendulum^31^; the exponent and clamping bounds in Eq. 25 were tuned empirically to keep simulated step times within physiologically plausible ranges^31–33^. In addition to step time, the generative model dynamically regulates step width based on a baseline anatomical width (*W*_*base*_) augmented by a lateral influence term. This term scales proportionately with the agent’s lateral velocity (|*v* _*y*_|) relative to its overall speed, where ϵ = 2. 2×10^−16^, directly mirroring compensatory base-of-support widening observed during human turning:

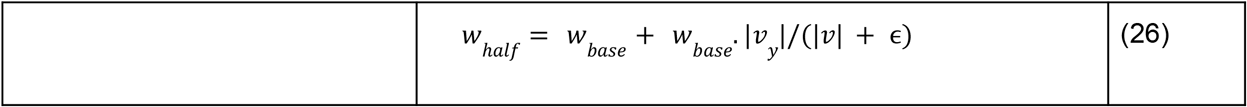

Eq. 26 gives the half-width used for foot placement. Step width is the measured perpendicular distance between consecutive footfalls, taken relative to the walking direction.

This resulting step time and dynamically modulated step width, combined with velocity, directly compute the distinct spatial-temporal gait parameters executed by the agent on the track, effectively capturing clinical observations. An example simulation showing step length and step width across a 90° turn is given in Fig. 1d and the LIP model is detailed in Fig. 1e.

#### Environment and the Reward Policy

The environment used in the model consists of a pathway which is followed by a turn and then another passage which leads to the target zone (Fig. 1a). The environment is designed in a way such that the track’s turning angle ϕ_*track*_ is the primary parameter which can be varied. The reward policy *r*(*t*) is designed to not only encourage reaching the goal but also to enforce safe turning kinematics. Whenever the agent successfully navigates the track and reaches the designated target zone, a terminal reward is given *r* =+ 2. Conversely, if the agent intersects the boundaries of the wall (collision), which depends on *r*_*body*_ denoted in Supplementary Table S1, the episode is terminated with a penalty *r* =− 1. For every safe step taken elsewhere on the track, the agent receives a small baseline positive reward *r* =+ 0. 01 to encourage continuous movement.

To facilitate optimal trajectory planning and speed regulation within the turn area itself, the baseline reward is augmented by three continuous shaping terms when the agent is inside the curve:

1. Lane Centering Bonus: A positive reward proportional to the agent’s distance from the nearer of the two track boundaries (*min_dist*), discouraging the agent from cutting close to either wall while navigating the curve (+ 0. 1. *min_dist*).
2. Alignment Bonus: A reward based on the dot product of the agent’s heading unit vector and the ideal tangential direction of the arc (+ 0. 05. *max*(0, *alignment*)), encouraging smooth and accurate steering.
3. Speed Penalty: A penalty directly proportional to the agent’s continuous velocity (− 0. 01. |*v*|), prompting the agent to decelerate to safely negotiate the turn radius.

Furthermore, upon successfully exiting the turn area, a minor alignment bonus (+ 0. 02. *max*(0, *alignment*)) is maintained to stabilize the agent’s heading and prevent post-turn collisions. These cumulative rewards are utilized to compute the temporal difference error δ(*t*) previously defined in Eq. 7, which iteratively trains the value function.

#### Simulating PD condition

To simulate the Parkinson’s Disease (PD) condition, the model represents dopamine deficiency by clamping the dopaminergic teaching signal — the temporal difference (TD) error δ(*t*), to a restricted upper limit, denoted as δ_*lim*_ ^26,34,35^. The PD category is further divided into two clinical subtypes: PD Freezers (PD-FOG) and PD non-Freezers (PD non-FOG)^36^. For the PD conditions, this capacity is strictly clamped:

PD non-FOG: δ = 0. 008, PD-FOG: δ _*lim*_= 0. 001, and δ _*med*_= 0, where δ_*med*_ is a parameter representing dopaminergic medication conditions in the model. Mathematically, this dopamine restriction is implemented as a conditional clamp on the TD-error during network training. If the PD condition is active (*pdcond* = 1) and the temporal difference error exceeds the defined limit, it is capped:

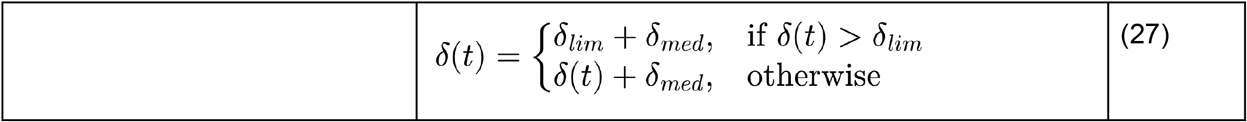

In our simulations δ_*med*_ is fixed at 0 for all conditions, since no pharmacological manipulationwas modeled here. The term is nonetheless built into Eq. 27 as an additive offset to the clamped TD-error, so that a positive δ_*med*_ value can partially restore the dopaminergic signal.This provides a direct mechanism for future work to simulate the effect of dopaminergic medication (e.g., levodopa) on turning-related freezing, without altering the model’s core architecture^37^.

To mathematically scale the action selection parameters between PD non-FOG and PD-FOG, a reference limit δ_*lim_ref*_ = 0. 01 is used to compute a disease severity multiplier called the FOG Factor:

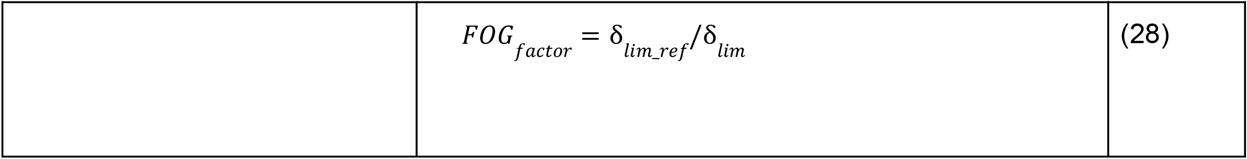

Rather than relying on static, pre-trained values, the model uniquely defines the disease manifestation as a dynamic interaction between the internal dopamine deficit (*FOG*_*factor*_) and the external environmental difficulty. The environmental difficulty is quantified by the angle ratio, defined as the track’s turning angle relative to a 90-degree corner:

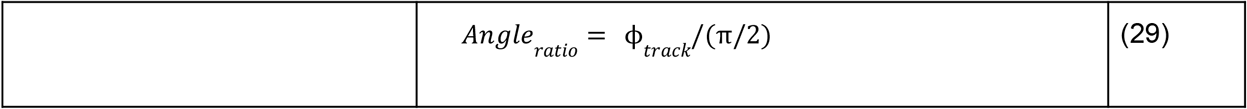

The main parameters that govern action selection — the Go amplitude *A*_*G*_, the No-go amplitude *A*_*N*_, and their respective slope parameters (λ_*go*_, λ _*nogo*_) begin at a baseline of *A*_*G*_ = 3 and *A* _*N*_= 3 with a base Go threshold θ _*go*_= 0, as mentioned in Table 3. For HC, these parameters remain unaltered, and the No-go threshold θ _*nogo*_remains strictly at 0.

However, when the PD condition is active, the direct pathway (Go) is suppressed and the indirect pathway (No-go) is pathologically amplified. This is modeled by scaling the baseline amplitudes and shifting the No-go activation threshold in direct proportion to both the disease severity multiplier and the track’s sharpness:

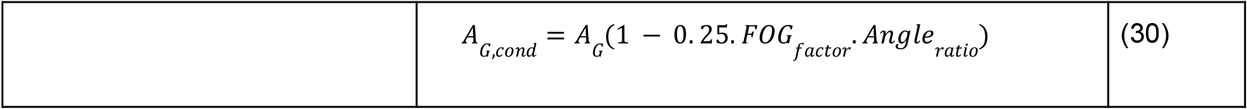

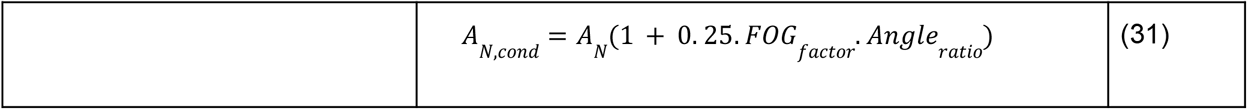

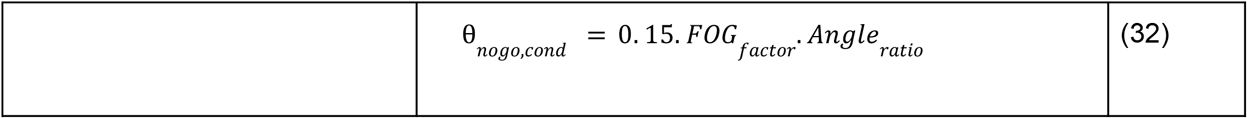

As the turn becomes sharper (increasing *Angle*_*ratio*_), a PD agent experiences an exacerbated drop in its movement-facilitating drive *A*_*G,cond*_ and a marked increase in its braking drive *A*_*N,cond*_ along with impaired estimation of value driven by clamping of the TD-error. Furthermore, the upward shift in the No-go threshold θ_*nogo,cond*_ means the braking pathway triggers earlier when the agent faces steep value gradients.

### Assessing the Model Performance

#### Value Function-Based Representation of Freezing and Slowness

The value function *V*(*t*) represents the agent’s estimated expectation of future rewards from specific spatial coordinates along the track. It is derived through temporal difference learning and it maps the optimal policy for efficient track traversal. The gradient of this value function Δ*V*(*t*) acts as a direct driving signal for the agent’s motor execution. In sharper turns, steep drops in the value gradient excite the No-go pathway and suppress the Go pathway. Consequently, clinical manifestations of bradykinesia and freezing are represented not just as low spatial values, but as pathologically amplified braking responses to these negative value gradients.

#### Spatial Gait Metrics: Step Length and Step Width

The model explicitly computes discrete spatial gait parameters using the LIP module. Step length is calculated at each motor update as the product of the agent’s continuous speed and the generated discrete step time *t*_*step*_. The step width is assessed by combining a baseline width with a dynamically scaling lateral influence term, driven by the ratio of lateral velocity to overall speed |*v* _*y*_|/(|*v*| + ϵ). These spatial metrics are averaged per episode over test trials to evaluate how they change across different turn severities and Parkinsonian conditions.

#### Temporal Gait Metrics: Step Time Variability

Step time variability is quantified to measure gait instability, a critical clinical marker of Parkinsonian freezing^38–40^. The model evaluates the temporal intervals of successive steps generated across the entire track to capture gait arrhythmia. The Coefficient of Variation (CV) was computed for each episode as the ratio of the standard deviation to the mean step time over all steps in test trials (*CV* = (σ/μ) × 100), and compared across 150 episodes per condition.

#### Kinematic Profiling: Turn Velocity and Smoothened Trajectories

The agent’s movement speed is profiled, particularly during the transition from straight walking to turning.

##### Turn Velocity

The model extracts the raw velocity magnitude (|*v*|) only when the agent’s spatial coordinates fall within the turn. This allows for direct comparison of steady-state turning speeds between Healthy Controls and PD agents.

##### Smoothened Velocity Profiles

To visualize turn-induced deceleration, the raw velocities of successful trials are extracted and processed using spline interpolation. This creates a normalized, smoothened continuous velocity curve across the length of the track, highlighting where braking pathways dominate the motor output.

#### Agent Trajectories and Success Metrics

To evaluate successful navigations, the agent’s Center of Mass (CoM) position coordinates (*x, y*) are tracked iteratively at each step. These coordinates are plotted sequentially to visualize the continuous spatial trajectory and assess lane-centering behavior in test trials. Simultaneously, we also track task success by evaluating boolean flags for each trial: a value of 1 is recorded if the agent reaches the target reward box, and a 0 is recorded if a wall collision occurs. These success metrics are aggregated over successive trial blocks to generate learning curves, quantifying how reliably the optimal turning policy is learnt. Success metrics are discussed in the Supplementary section and are visualized in Supplementary Fig. S2.

#### Parameter Space Analysis: Velocity and Value Heatmaps

Dopaminergic dysfunction in PD is mainly thought to impair two distinct RL-related processes. One is the encoding of reward-prediction error itself, and second is the regulation of exploratory behavior during action selection. Previous basal ganglia models have simulated the former by constraining the TD-error to reflect the degraded value signal seen in PD gait^18^, while separate work has proposed the basal ganglia, particularly the indirect pathway, as a substrate for exploration, with dopamine loss linked to reduced exploratory drive^34,41^. To investigate this interaction between dopamine deficiency and exploratory behavior, the model’s outputs are evaluated across a two-dimensional parameter space mapping the dopaminergic TD-error limit δ against the exploration amplitude (*Ae*) using the trajectories generated during test trials.

Value Function Heatmaps: The expected mean state values computed by the Critic network are mapped across the δ_*lim*_ − *Ae*_*eff*_ parameter space (Fig. 2c). This demonstrates how the internal neural representation of the track’s spatial reward degrades or shifts under the combined influence of clamped dopamine limits δ_*lim*_ and varying exploratory gains *Ae*_*eff*_.

**Figure 2.**
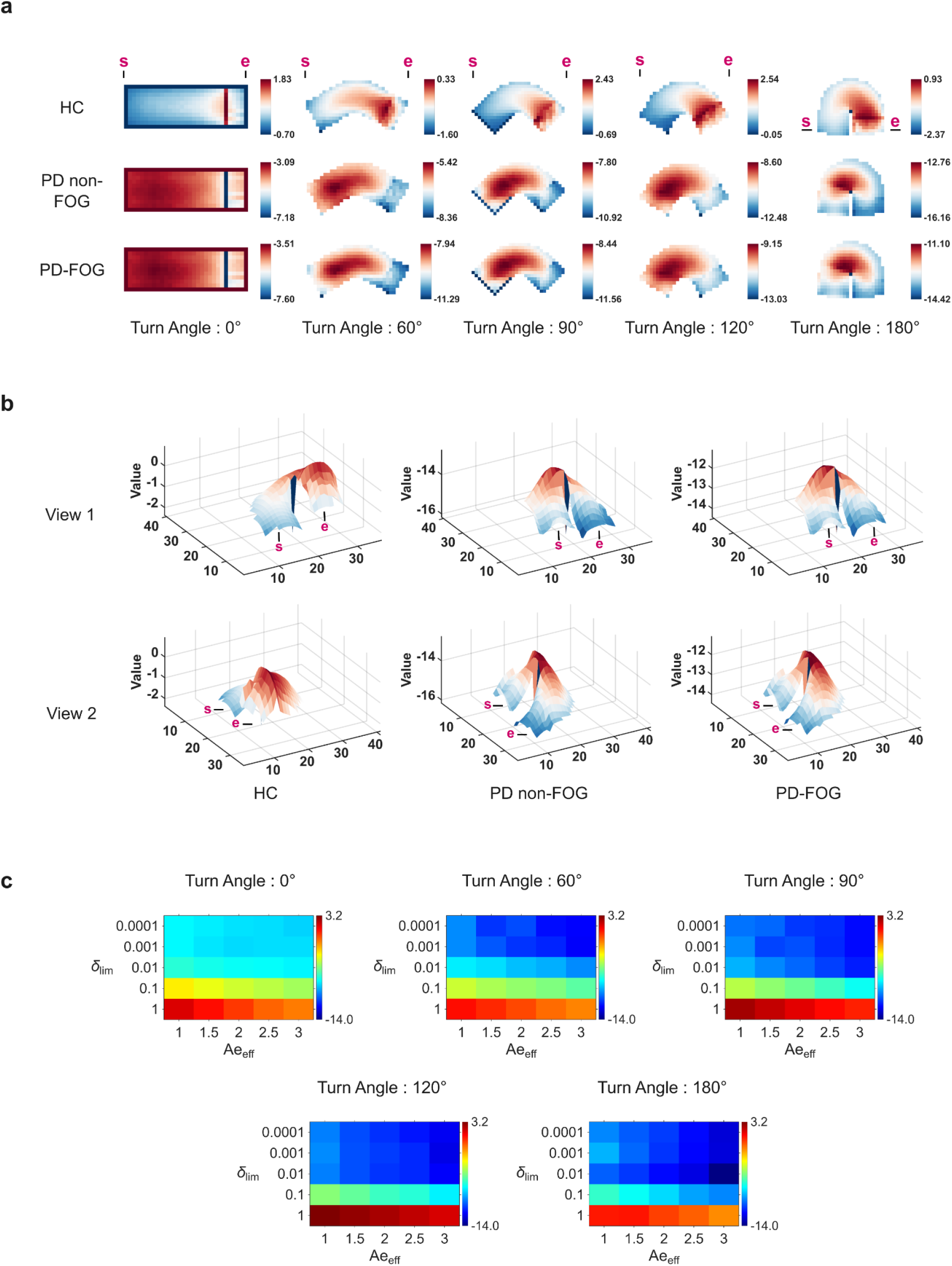
**Value Function Representations and Parameter Space Heatmaps:** (a) Two-dimensional heat maps of the Critic’s learned value function *V*(*t*), plotted over the (x, y) positions of the track for Healthy Controls (HC), PD non-FOG, and PD-FOG, at 0°, 60°, 90°, 120°, and 180° turn angles. Color scales are independent per panel. Each map shows the value the trained Critic assigns to every position on the track, using the final network weights from each of the 150 episodes, averaged across seeds, with warmer colors (red) indicating higher estimated state value and cooler colors (blue) indicating lower estimated value. ‘s’ marks the start of the track and ‘e’ marks the end of the track. HC shows a smooth increase in the value function, with the highest values concentrated near the reward zone. In contrast, both PD groups show an overall depressed value landscape, with high values near the start of the track and lower values near the reward box. (b) Three-dimensional surface plots of *V*(*t*) over the (x, y) positions of the 180° track for HC (left), PD non-FOG (center), and PD-FOG (right), shown from two viewing angles (View 1, View 2). Color scales are independent per panel. ‘s’ denotes the start of the track, while ‘e’ denotes the end. HC shows a steady increase in value up to the reward box, while both the PD conditions show a topographic dip in the turn, with lower values near the reward box. (c) Heatmaps of mean *V*(*t*) along the agent’s test trajectories, mapped across the TD-error clamp limit (δ_*lim*_ ; y-axis, 0.0001–1) and the exploration amplitude (*Ae*_*eff*_ ; x-axis, 1–3), across all turn angles. Warmer colors indicate higher estimated value; color scales are directly comparable across angles. Mean value generally increases with δ_*lim*_ at every turn angle, from the lowest values at δ_*lim*_ ≤ 0.01 to highest at δ_*lim*_ = 1, while *Ae*_*eff*_ has a comparatively small effect.

Velocity Heatmaps: Similarly, these heatmaps visualize the average turn velocity of the agent across the δ_*lim*_ − *Ae*_*eff*_ grid, highlighting the specific parameter boundaries where FOG might occur. Parameter space analysis for velocity is discussed in the Supplementary section, with Supplementary Fig. S3 detailing the heatmaps.

## RESULTS

### Value Function Representations in Turning Environments

To understand how the value function evolves over different track turning angles, we analyzed the average value function at a given position for all possible state vectors. We simulated the agent traversing tracks with different turning angles (0°, 60°, 90°, 120°, and 180°). The average 2D value function for 150 episodes of the agent is visualized in Fig. 2a. Healthy Controls successfully formed a value function capable of navigating complex tracks. The expected reward progressively increased near the turning area and the final reward box, while sharply dropping along the boundaries. This reflects a robust, learned policy to seek the goal while avoiding wall collisions.

In contrast, training the model with a clamped TD-error to simulate PD fundamentally altered this spatial representation in PD non-FOG and PD-FOG. The value progressively dropped from the turn apex to the reward box. There was an abnormally increased value at the agent’s starting location in PD, suggesting they would prefer to remain at the start rather than move forward.

This altered spatial topography is detailed in Fig. 2b, which provides 2 views for 3D surface visualization of the value function for the 180° turn. While the HC’s value steadily climbs toward the goal, both PD non-FOG and PD-FOG exhibit a dip near the reward box after the turn. This mirrors a similar pattern from our earlier study modeling doorway freezing in PD^18^, where we showed a decrease in the value function close to the doorway that scaled with its narrowness.

All these results suggest that training the model with a clamped TD-error degrades the spatial reward representation, creating a “value canyon” that the agent struggles to cross. This degraded value function provides a mechanistic explanation for the behavioral results seen in experiments, clarifying why turn freezing probability is so high when turning sharpness increases. A visual representation of the turning strategies used by HC, PD non-FOG and PD-FOG for ∼10 timesteps under their respective conditions is shown in Supplementary Fig. S1.

### Parameter Space Analysis: Value Heatmaps

To understand how dopaminergic depletion and exploration amplitude jointly shape the agent’s value estimates, we evaluated mean *V*(*t*) across a 2D grid mapping the TD-error clamp limit δ_*lim*_ against the exploration amplitude *Ae*_*eff*_ (Fig. 2c). These values were computed by averaging *V*(*t*) across all steps of all testing trials. At all turn angles, δ_*lim*_ was the dominant determinant of value. A sharp decrease in value is observable when δ_*lim*_ is decreased, which is more drastic for sharper turns. The highest values for all turn angles are seen at δ_*lim*_ = 1. In comparison, *Ae*_*eff*_ had a smaller effect. At δ_*lim*_ = 1, higher *Ae*_*eff*_ reduced value at all angles, more pronounced for shallower turns, consistent with excessive exploration during learning being detrimental when the path is simple. However, this drop in value (at δ_*lim*_ = 1) is not so steep for 180°, suggesting the need for a slightly higher exploration rate for sharper turns.

### Agent Trajectories

To visualize how the internal value representation translates into physical movement, we extracted the two-dimensional spatial trajectories of the agents as they navigated the track, shown in Fig. 3a. The successful trials are plotted in bold colors, while failed trials (e.g., wall collisions or aborted movements) are plotted as faint grey traces. HC shows high success rates (Supplementary Fig. S2) as well as deterministic trajectories. Their spatial trajectories were smoothly executed and tightly clustered, indicating consistent path optimization with precise motor control.

**Figure 3.**
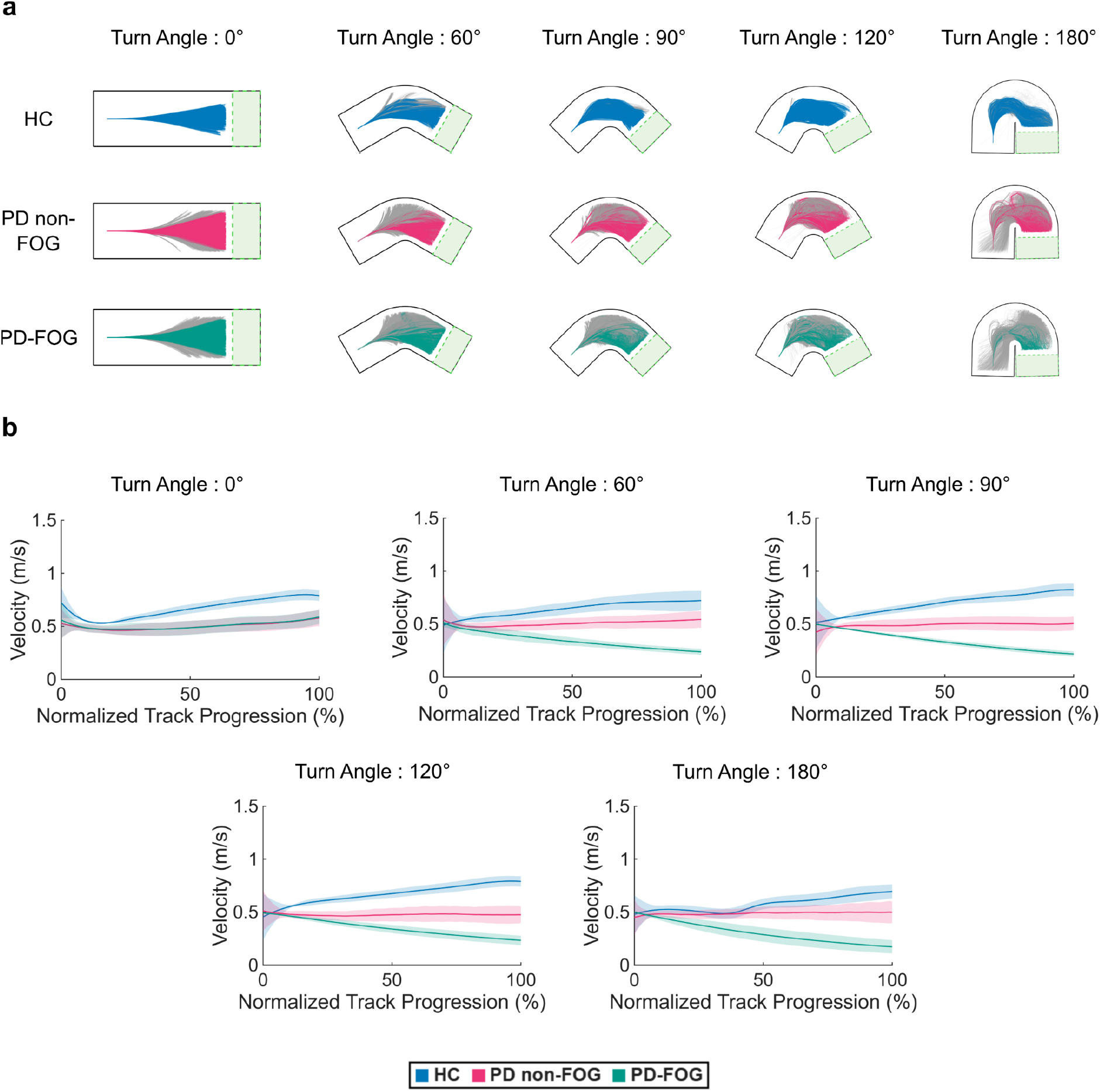
**Agent Trajectories and Velocity Profiles:** (a) Overhead view of individual agent trajectories overlaid on track geometry across turn angles (0°, 60°, 90°, 120°, and 180°) for Healthy Controls (HC; top row, blue), PD non-FOG (middle row, pink), and PD-FOG (bottom row, teal). Colored traces indicate successful test trials reaching the reward zone (green dashed box); faint grey traces indicate failed test trials that did not reach the goal. HC trajectories form a tight bundle that closely hugs the inner boundary of the turn. In both PD groups, the proportion of failed (grey) trials increases with turn angle, showing more hesitation and erratic steps in PD-FOG, consistent with the reduced success rate shown in Supplementary Fig. S2. (b) Mean velocity (± standard deviation, shaded band) across successful test trials as a function of normalized track progression (0% = track entry, 100% = reward area) for HC, PD non-FOG, and PD-FOG, shown for each turn angle. All conditions begin from a common velocity (0.5 m/s) at track entry. At 0°, HC accelerates, whereas both PD groups maintain a lower velocity. From 60° onwards, PD-FOG shows a monotonic velocity decline across the turn, diverging from both HC (which moderately accelerates through the turn) and PD non-FOG (which remains roughly flat), consistent with an in-turn deceleration rather than a fixed offset in baseline speed.

In contrast, the simulated dopamine depletion in PD non-FOG and PD-FOG resulted in a marked deterioration in the ability to turn, which scaled directly with the turn angle. In PD non-FOG, introducing a curve caused widening of the trajectory spread, indicating spatial instability. As the turn angle sharpened, the proportion of failed trials increased significantly, with agents frequently colliding with the boundaries near the turn’s apex.

This spatial impairment was most extreme in the PD-FOG. While the agent managed a limited number of successful runs at shallow angles, sharp turn angles (≥90°) triggered a near-total breakdown in motor execution. The plots reveal dense clusters of grey failure traces heavily concentrated at the initiation and apex of the turn, with very few trajectories surviving to the reward box.

### Velocity Profiles in Turn Execution

To quantify the kinematic impact of the altered value representations, we analyzed the average velocity profiles of the agents as they executed the turns. Fig. 3b displays the smoothened velocity of successful test trials, normalized from track entry (0%) to exit (100%). Raw velocity trajectories from successful test trials were resampled onto a standardized 100-point grid via cubic spline interpolation and averaged across episodes. HC maintained fluid motor execution and high velocity across all turn angles, exiting at ∼0.7–0.8 m/s. This reflects an unimpaired basal ganglia circuit capable of smoothly linking motor sequences to match a healthy value gradient.

In contrast, the simulated dopamine depletion in the PD models produced distinct, condition-specific kinematic impairments. PD non-FOG’s velocity remained relatively flat and significantly lower than the HC agent (∼0.5 m/s) across all angles. While slower and more cautious, among successful trials it maintained this steady velocity, allowing itself to successfully complete the turns without halting.

PD-FOG demonstrated a severe, progressive deceleration that was dynamically linked to the geometry of the track. While its velocity remained stable on the straight (0°) track, the introduction of any curve (≥60°) caused a continuous degradation of speed as the agent progressed deeper into the turn. At sharper angles, the agent’s velocity plummeted to nearly 0.2 m/s by the end of the turn.

### Gait Metrics

We analyzed the gait metrics between HC, PD non-FOG and PD-FOG during the testing phase. The simulated discrete spatial and temporal parameters explicitly capture the hallmark clinical symptoms of Parkinsonian gait across varying turns as shown in Fig. 4. Across all gait metrics, statistical differences were evaluated using two-way mixed-design ANOVA to test for main effects of condition and turn angle, as well as their interaction. All post-hoc pairwise comparisons were Bonferroni-corrected for multiple comparisons within each family (3 group comparisons per angle; 10 angle comparisons per group). Mauchly’s test indicated that the assumption of sphericity was violated for all four gait metrics (all p < 0.001); degrees of freedom were corrected using Greenhouse-Geisser estimates where ε ≤ 0. 75, and Huynh-Feldt estimates where ε > 0. 75, following standard convention^42^.

**Figure 4.**
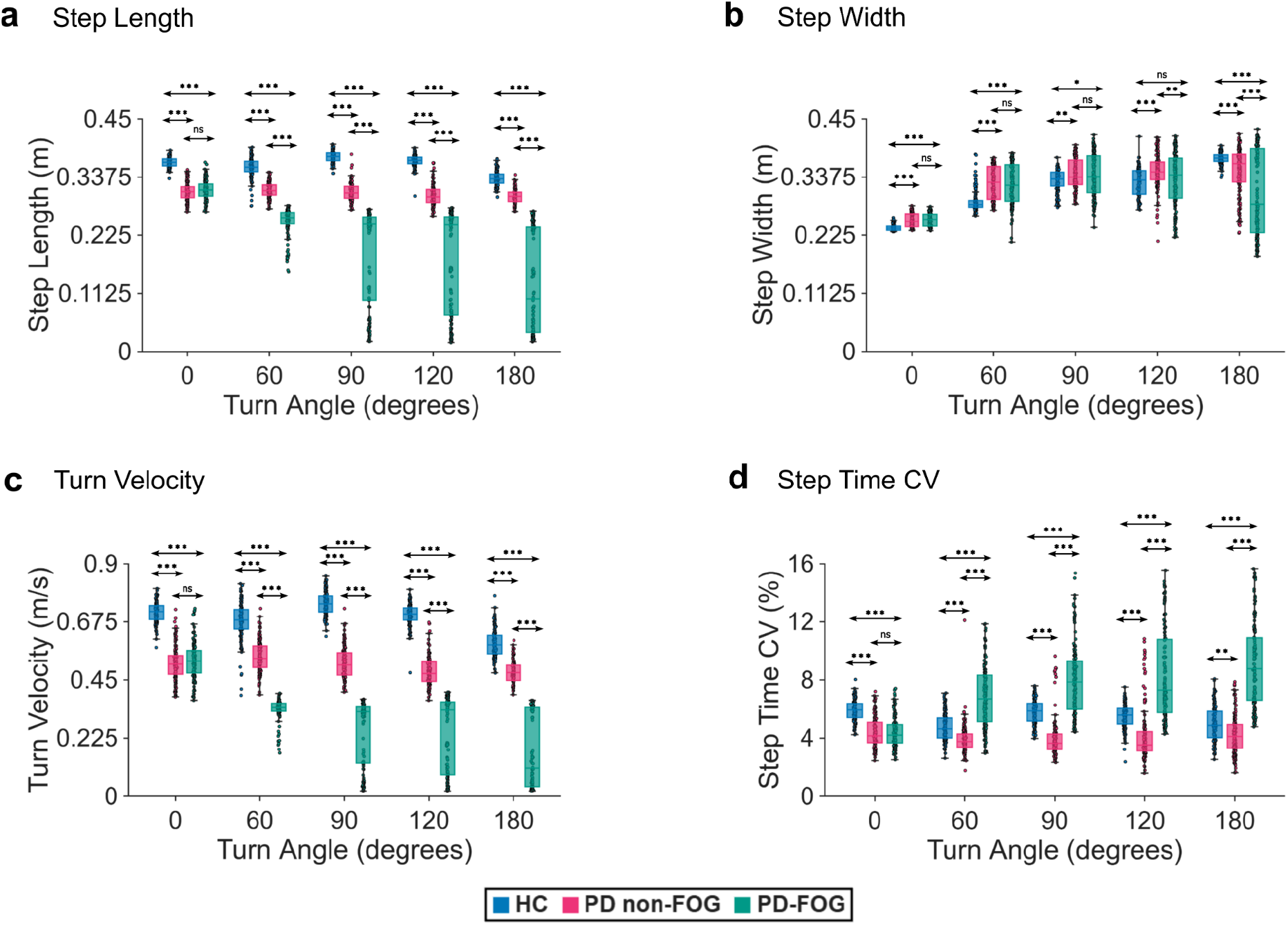
**Gait Metrics Across Turn Angles:** Gait parameters for Healthy Controls (HC), PD non-FOG and PD-FOG across all turn angles (0°, 60°, 90°, 120°, 180°). (a) Step length (m). (b) Step width (m). (c) Turn velocity (m/s). (d) Step time coefficient of variation (CV, %). Box plots show the median and interquartile range across 150 episodes (test trials) per condition and turn angle. Overlaid scatter points represent the mean value of each individual episode. Asterisks indicate Bonferroni corrected pairwise comparisons for all three group pairs (HC vs. PD non-FOG, HC vs. PD-FOG, PD non-FOG vs. PD-FOG) at each turn angle (*p < 0.05, **p < 0.01, ***p < 0.001, ns = not significant). Overall two-way mixed-design ANOVA statistics (main effects of condition and angle, and condition × angle interaction) and within-condition across-angle post-hoc comparisons are reported in the main text.

#### Step Length

Significant main effects of turn angle (F_(2.88,1286.14)_ = 155.29, p < 0.001, ηp^2^ = 0.26, Greenhouse-Geisser corrected) and condition (F_(2,447)_ = 2057.5, p < 0.001, ηp^2^ = 0.90), as well as a significant Condition × Angle interaction (F_(5.75,1286.14)_ = 102.94, p < 0.001, ηp^2^ = 0.32, Greenhouse-Geisser corrected) were revealed for step length, and pairwise comparisons at each angle are shown in Fig. 4a. At 0°, HC differed significantly from both PD groups (HC vs. PD-FOG: p < 0.001; HC vs. PD non-FOG: p < 0.001), while the two PD groups did not differ from each other (p = 0.168), specifying that the FOG-specific deficit is not present at baseline. From 60° onward, all three pairwise comparisons reached significance at every angle (60°, 90°, 120°, 180°; all p < 0.001), indicating that PD-FOG separates from PD non-FOG as soon as turning begins.

Comparisons across angles, analyzed separately within each condition, failed to find any significant difference in PD non-FOG step length at any angle versus any other (all p > 0.15). This suggests a stable shortening relative to HC rather than a progressive decline. HC showed a small decrease from 0° to 60° (p < 0.001), a recovery from 60° to 90° (p < 0.001), and a significant decline at 180° relative to every other angle (all p < 0.001). In PD-FOG, step length reduced significantly from 0° to every subsequent angle (60°, 90°, 120°, 180°; all p < 0.001), continuing to decline significantly with each further angle step (0°→60°, 60°→90°, 60°→120°, 90°→180°, 120°→180°; all p < 0.001) with the only exception of 90° to 120° which failed to reach significance (p = 0.20), showing a brief plateau followed by further decline toward the sharpest angle. This pattern matches the shuffling, short-stepped gait that precedes clinical freezing in PD-FOG. This is largely consistent with Bhatt et al.^10^ who found that both PD groups tended to walk with shorter step lengths compared to HC, and that step length decreased systematically with increasing sharpness of the turn for all groups (main effect of angle, F_(3,81)_ = 154.38, p < 0.001). Our model reproduces this reduced step length in both PD conditions relative to HC, and additionally captures a deteriorating trend as a function of turning angle specific for PD-FOG.

#### Step Width

We found significant main effects of turn angle (F_(2.88,1285.36)_ = 533.2, p < 0.001, ηp^2^ = 0.54, Greenhouse-Geisser corrected) and condition (F_(2,447)_ = 27.34, p < 0.001, ηp^2^ = 0.11), plus a significant Condition × Angle interaction (F_(5.75,1285.36)_ = 44.73, p < 0.001, ηp^2^ = 0.17, Greenhouse-Geisser corrected) for step width. Pairwise comparisons at each angle are shown in Fig. 4b. At baseline, HC already differed from both PD groups, which reflects a compensatory base-of-support widening from the start. HC’s step width was significantly narrower at 0° (HC vs. both PD groups: p < 0.001), but the two PD groups did not differ from each other (p = 0.924). This HC-narrower pattern persisted till 60° (both p < 0.001) and at 90° (HC vs. PD-FOG: p = 0.026; HC vs. PD non-FOG: p = 0.0015), with the two PD groups still not differing from each other at either angle (both p = 1). By 120°, HC was no longer significantly narrower than PD-FOG (p = 0.38), though it remained significantly narrower than PD non-FOG (p < 0.001). The two PD groups diverged at 120°, with PD-FOG becoming significantly smaller than PD non-FOG (p = 0.0023), and at 180° (p < 0.001).

Comparisons across angles showed that HC widened its step progressively across almost every turn angle (0°→60°, 0°→90°, 0°→120°, 0°→180°, 60°→90°, 60°→180°, and 120°→180°: all p < 0.001). This indicates that HC continues widening its stance as turns get sharper. PD non-FOG widened significantly from 0° to all angles (all p < 0.001) and continued widening further from 60° to both 90° and 120° (both p < 0.001), but showed no further significant change among 90°, 120°, and 180° (all p = 1), showing that it first widens its base and then holds it from 90° onward. PD-FOG initially matched this strategy, widening significantly from 0° to 120° (all p < 0.001) and plateauing between 90° and 120° (p = 1), but then reversed: step width dropped significantly from both 90° and 120° to 180° (both p < 0.001), and also from 60° to 180° (p = 0.015). PD-FOG’s step width at 180° remained significantly above its own 0° baseline (p < 0.001) — indicating a partial collapse of the compensatory response.

This early, group-wide compensatory widening that becomes selectively unsustainable in PD-FOG at the sharpest turn parallels Bhatt et al.^10^, who reported a significant Group × Angle interaction for step width (F_(6,81)_ = 3.746, p < 0.0025) and found that PD-FOG patients failed to widen their base of support at sharp turning angles in contrast to HC and PD non-FOG, who both sustained widening. Our model captures this selective breakdown where PD-FOG widens with HC and PD non-FOG for moderate turn angles, then reverses sharply at 180°, falling significantly below both its own 90°/120° peak and both comparison groups (all p < 0.001).

#### Turn Velocity

The velocity results clearly show the severity of clinical bradykinesia and freezing (Fig. 4c). Significant main effects of turn angle (F_(3.61,1612.47)_ = 265.93, p < 0.001, ηp^2^ = 0.37, Huynh-Feldt corrected) and condition (F_(2,447)_ = 4428.9, p < 0.001, ηp^2^ = 0.95), as well as Condition × Angle interaction (F_(7.21,1612.47)_ = 129.87, p < 0.001, ηp^2^ = 0.37, Huynh-Feldt corrected) were reported on turn velocity. At 0°, HC was already significantly faster than both PD groups (both p < 0.001), while PD-FOG and PD non-FOG did not yet differ from each other (p = 0.44). From 60° onward, all three pairwise comparisons were significant at every angle (HC vs. PD-FOG, HC vs. PD non-FOG, and PD-FOG vs. PD non-FOG: all p < 0.001 at 60°, 90°, 120°, and 180°), with PD-FOG consistently the slowest of all.

Comparisons across angles revealed distinct within-group trajectories. HC showed a small dip in turning velocity at 60° (p < 0.001) followed by a rise back above baseline at 90° (p < 0.001) with no net change between 0° and 120° (p = 1), before a much larger decline at 180° relative to every other angle (all p < 0.001). This indicates that even healthy agents slow down substantially when turns get sharper. In PD non-FOG, velocity rose slightly above baseline at 60° (p < 0.001), then declined significantly relative to baseline by 120° and 180° (120°: p = 0.014; 180°: p = 0.002), plateauing between 120° and 180° (p = 1) — a small, delayed decline rather than stable bradykinesia. PD-FOG, in contrast, declined immediately and severely: velocity dropped significantly from 0° to every subsequent angle, with the drop already present by 60° (p < 0.001). It continued declining from 60° to 90° and 60° to 120° (both p < 0.001), plateaued between 90° and 120° (p = 1), and finally declined significantly to 180° (both relative to 90° and 120°, p < 0.001).

This severe, early-onset, geometry-dependent deceleration in PD-FOG contrasted with HC’s late-onset slowing and PD non-FOG’s smaller, delayed decline — captures the motor arrests observed in turn-induced freezing, and is broadly consistent with Bhatt et al.^10^, who reported a significant Group × Angle interaction for velocity (F_(3,54)_ = 4.915, p < 0.005), with freezers showing a progressive decrease in velocity from straight walking through all turning angles. It is also consistent with Gilat et al.^20^, who found that PD patients with FOG had significantly longer scaled maximum footstep latencies than non-freezers specifically in turns in a virtual-reality paradigm (p = 0.028), while non-freezers showed no significant difference between turning and walking latencies (p = 0.506), paralleling our finding that freezing-related slowing in PD-FOG emerges steeply with turning demand rather than existing as a fixed baseline trait.

#### Step Time CV

As a primary indicator of gait arrhythmia, step time CV captures the irregular, hesitant stepping characteristic of freezing episodes (Fig. 4d). Significant main effects of turn angle (F_(3.51,1569.97)_ = 47.55, p < 0.001, ηp^2^ = 0.10, Huynh-Feldt corrected) and condition (F_(2,447)_ = 639.16, p < 0.001, ηp^2^ = 0.74), as well as Condition × Angle interaction (F_(7.02,1569.97)_ = 74.01, p < 0.001, ηp^2^ = 0.25, Huynh-Feldt corrected) were revealed for step time CV. At 0°, HC’s CV was significantly higher than both PD groups (HC vs. PD-FOG: p < 0.001; HC vs. PD non-FOG: p < 0.001), while the two PD groups did not differ from each other (p = 1). The comparison with PD-FOG reverses direction once turning begins: from 60° onward, PD-FOG becomes significantly more arrhythmic than both HC and PD non-FOG at every subsequent angle (all p < 0.001).

Notably, HC’s CV remained significantly higher than PD non-FOG’s at every angle tested (60°, 90°, 120°, 180°, all p < 0.01). This is consistent with the sigmoid nonlinearity governing speed control (Eqs. 9, 10): across all turn angles tested, HC’s Δ*V*(*t*) remained more tightly concentrated near zero than PD non-FOG’s, which showed a wider distribution at every angle. Sigmoid is most sensitive to inputs near its center and increasingly insensitive to inputs away from the center. Hence, HC’s narrower signal passed through largely undamped into speed and step timing, while most of PD non-FOG’s wider inputs fell into the sigmoid’s saturated range, compressing the variability. At 180°, this compression was further compounded by θ_*nogo, cond*_, adding a second source of compression on top of the distribution-width effect present at every angle.

Comparisons across angles revealed that PD non-FOG’s step time CV was largely flat, with only one significant dip from 0° to 60° (p < 0.01) and no additional significant differences between any other pair of angles (all p = 1 or p > 0.05). In HC, CV decreased from 0° to 60° (p < 0.001) and from 0° to 180° (p < 0.001), rose again between 60° and 90° (p < 0.001) and 60° and 120° (p < 0.01), then declined once more from 90° to 180° (p < 0.001). In contrast, PD-FOG’s variability rose significantly and progressively relative to its own 0° baseline at every subsequent angle (60°, 90°, 120°, 180°, all p < 0.001), with significant increases also from 60° to 90° and 60° to 120° (both p < 0.001). This rise briefly plateaued between 90° and 120° (p = 1), but resumed with a further significant increase from both 90° and 120° to 180° (both p < 0.01), indicating that the elevated variability in PD-FOG continues to worsen at the sharpest turn.

This progressive rise in step time CV in PD-FOG mirrors the irregular, hesitant stepping sequences characteristic of freezing of gait triggered by complex motor tasks, and matches Bhatt et al.^10^, who found a significant Group × Angle interaction for step time variability (F_(6,81)_ = 3.791, p < 0.005), with freezers showing significantly greater variability than healthy controls specifically at the 120° and 180° turns. It also aligns directionally with Gilat et al.^20^, who reported that PD patients with FOG had significantly higher step time CV than non-freezers both during turning (16.5% vs. 9.6%, p < 0.01) and during simple walking (15.7% vs. 9.3%, p < 0.01) in the virtual-reality gait task. This suggests that our model captures the clinically observed elevation in temporal variability among freezers, which scales with turn sharpness.

## DISCUSSION

We successfully captured turn freezing dynamics in PD using a cortico-basal ganglia-inspired model that captures the deficits in processing value function over space. Our model captured key gait characteristics, including turn velocity and step length reduction as turn severity rose and step time arrhythmicity, along with a breakdown of compensatory step width widening in PD-FOG at the sharpest turn angle. We leveraged the idea that the clamped levels of temporal difference error, a model surrogate for dopamine reduction, could lead to changes in the value function computation over the track, an approach which explained narrow doorway freezing and freezing during cognitive load in our previous studies^17,18^. Similar to our observations on modeling doorway freezing^18^, there was a drop in the value function near the point of the turn that influenced the Value Difference Δ*V*(*t*) affecting action selection operating via the Go/Explore/No-go (GEN) dynamics. This effect was compounded by δ_*lim*_ ‘s rescaling of the Go/No-go gains and the θ_*nogo,cond*_ at every action-selection step. Together, they provide a potential explanation as to why PD non-FOG sustains lower speeds without progressive deceleration, while PD-FOG shows excessive No-go signaling, culminating in severe gait arrhythmicity and motor arrests.

In the many subtypes of freezing behavior, turn freezing is one of the most prominent contexts triggering motor arrests^15^. Unlike narrow doorway/passage freezing, turn freezing scales directly in difficulty with turn angle^10^ due to asymmetric motor and base-of-support adjustments^43^. Hence, we found that the model training required introduction of scaling parameters to retune the agent’s strategy for a given turn sharpness. In the PD conditions, this was done by combining the environment sharpness (*Angle*_*ratio*_) with the FOG factor (*FOG*_*factor*_) derived from the clamped TD-error limit (δ_*lim*_), leading to hyperactivation of the indirect pathway (*A*_*N,cond*_ and θ_*nogo,cond*_), while simultaneously suppressing the direct pathway drive (*A*_*G,cond*_). Notably, this Go/No-go asymmetry was absent in the HC model, which retained the baseline gains (*A*_*G*_, *A*_*N*_ and θ_*nogo*_ = 0) with no TD-error clamping at all, isolating dopaminergic deficit-induced freeze.

Our model also required scaling some of the gain parameters of the GEN with the turn angle applied uniformly across all three groups. This included the exploration amplitude (*Ae*_*eff*_) and its decay variance (σ), the steering probe angle (δ_*steer*_), the steering gain (*K*_*steer*_), and the turning rate (*K*_*turn*_), all of which scaled dynamically with the *Angle*_*ratio*_. Navigating sharper turns requires probing headings further ahead, applying stronger steering corrections once a value gradient is detected, and exploring longer during training. To prevent erratic oscillations, the steering signal was dampened by 70% upon exiting the turn, which is analogous to frontostriatal motor set-shifting needed to flexibly disengage a turning motor program and re-stabilize again for straight paths. Only when these gains were scaled to match the sharpness of the turn could HC produce the smooth turning seen in Fig. 3b. Adding *FOG*_*factor*_ - driven Go/No-go dynamics to this geometric scaling causes turns that HC navigates smoothly to pathologically suppress action drive in PD groups.

The critical parameter differentiating HC from both PD groups was the TD-error clamp δ_*lim*_, our model’s proxy for reduced phasic dopaminergic signaling from the SNc^26^. In HC, the TD-error was left unclamped, allowing the Critic to freely update its value estimates and produce the smooth, monotonically increasing value gradient toward the goal seen in Fig. 2a. In PD groups, clamping this signal degraded reward prediction errors, creating the “value canyon” near the turn apex and reward area. This is consistent with the classical framing of dopamine as the teaching signal underlying corticostriatal plasticity^44^ — when phasic dopamine bursts are blunted, as in PD, the striatum can no longer accurately update its estimate of state value, particularly in regions of the environment (like a sharp turn) that demand rapid, fine-grained value updates. The magnitude of this clamp separated PD non-FOG from PD-FOG (δ_*lim*_ = 0.008 vs. 0.001), producing a larger *FOG*_*factor*_ in the PD-FOG condition. In basal ganglia terms, this maps onto the classical direct (striatonigral, D1-mediated) versus indirect (striatopallidal, D2-mediated) pathway framework. Dopamine depletion is known to preferentially impair D1-dependent Go signaling while simultaneously disinhibiting D2-dependent No-go signaling^26^, increasing net inhibitory output from the basal ganglia to the thalamocortical motor circuit. This asymmetric *A*_*G, cond*_ and *A*_*N, cond*_ scaling computationally mirrors the pathway imbalance. The fact that severe freezing emerged only at sharper turns suggests that FOG occurs when task demand and dopamine loss combine multiplicatively to cross a threshold, which is consistent with clinical observations that FOG is episodically triggered rather than continuous^6^.

Our model contains some limitations. First, we represent dopaminergic depletion using a single clamped parameter (δ_*lim*_) on the TD-error, which is a simplification of the complex neurochemical changes occurring in PD. Furthermore, while we held the medication offset (δ_*med*_) at zero, future work could vary δ_*med*_ to simulate dose-dependent drug effects on turning-related FOG^6^. Second, several gain parameters in the GEN block (e.g., *Ae*_*eff*_, *K*_*steer*_, *K*_*turn*_) were manually scaled with turn angle to allow the model to learn sharper turns. Future work can focus on how such adaptive gain modulation could emerge endogenously from the learning process itself. Third, our leg model uses a simplified Linear Inverted Pendulum representation and does not capture detailed joint-level or muscular dynamics involved in real human gait. Finally, we tested a limited set of discrete turn angles in isolation, whereas real-world freezing often results from turning combined with other factors such as dual-tasking or fatigue, which are not modeled here. Despite these simplifications, the model successfully captures the key kinematic and spatial features of turn freezing observed clinically, and provides a useful framework for understanding the computational basis of FOG.

Overall, our work suggests a generalized framework for approaching disorders which deal with the evaluation of space. The deficits that arise due to the clamping of the dopaminergic teaching signal are not specific to any one spatial context — the same mechanism that explained narrow doorway freezing^18^ and cognitive-load-induced freezing^17^ also reproduces the kinematic hallmarks of turn freezing once the model is extended with the turn-specific geometric scaling described above. This suggests that FOG may share a common underlying computational deficit: a degraded internal value representation of the space the patient must navigate, rather than distinct, context-specific pathologies. Under this framework, the specific triggers of freezing such as doorways, turns, dual-tasking, can be understood as different environmental demands that expose the same core impairment, where it is the steepness or discontinuity of the local value gradient, rather than the nature of the trigger itself, that ultimately determines whether a motor arrest occurs.

## Code and Data Availability Statement

The source code for the computational model, along with the data generation scripts and plotting routines will be publicly available on ModelDB upon acceptance of the manuscript.

## Author Contributions

Conceptualization: Neermita Bhattacharya, Sandeep Sathyanandan Nair, V. Srinivasa Chakravarthy, Vignesh Muralidharan; Methodology: Neermita Bhattacharya, Sandeep Sathyanandan Nair, V. Srinivasa Chakravarthy, Vignesh Muralidharan; Model Development: Neermita Bhattacharya and Sandeep Sathyanandan Nair; Formal analysis: Neermita Bhattacharya and Sandeep Sathyanandan Nair; Visualization: Neermita Bhattacharya; Supervision: Vignesh Muralidharan; Writing – original draft: Neermita Bhattacharya, Sandeep Sathyanandan Nair, Vignesh Muralidharan; Writing – review & editing: Neermita Bhattacharya, Sandeep Sathyanandan Nair, V. Srinivasa Chakravarthy, Vignesh Muralidharan.

## Funding

Vignesh Muralidharan would like to acknowledge the research funds from Indian Institute of Technology Jodhpur (IIT Jodhpur) [No specific grant number is associated with this funding].

## Declaration of competing interests

The authors declare no competing interests.

## Acknowledgements

We thank the Indian Institute of Technology Jodhpur for both the infrastructure and computational resources. We also sincerely thank Arpit Dilipbhai Shrimankar for his assistance in running the HPC simulations.

The authors used AI assistance tools (Google Gemini 3.5 Pro and Claude Sonnet 5.5) for limited support in code refinement and language editing of the manuscript. The tools did not contribute to the model development, data analysis, interpretation of results, or scientific conclusions. The authors remain fully accountable for originality, accuracy, and integrity of the content.

## Notes

### Competing Interest Statement

The authors have declared no competing interest.

