## Supplementary Information for "Impaired value estimation over space explains turn-induced freezing of gait in Parkinson’s disease: A cortico-basal ganglia-inspired reinforcement learning model"

**Supplementary Table S1: Leg Model Parameters (LIP / Gait Generation)** — biomechanical constants governing the Linear Inverted Pendulum model used to generate stepping kinematics.

| **Parameter** | **Symbol** | **Value** | **Description** |
|---|---|---|---|
| Gravitational Constant | $g$ | $9.81 m/s^2$ | Used to derive LIP natural frequency $\omega = \sqrt{g/h}$. |
| CoM Height | $h_0$ | $0.9 m$ | Baseline anatomical Center of Mass height. |
| Step Time Speed Exponent | — | $0.36$ | Exponent in Eq. (25). |
| Step Time Clamp Bounds | — | $0.25 s, 0.9 s$ | Min/max in Eq. (25). |
| Body Radius | $r_{body}$ | $0.25 m$ | Physical footprint for boundary-collision detection. |
| Base Half-Period | $T_{base}$ | $\pi/2\omega$ | Base half-period of the LIP model in (Eq. 25) |
| Baseline Half Step-Width | $w_{base}$ | $0.12 m$ | Anatomical baseline half-width in Eq. (26). |
| Division Safeguard | $\epsilon$ | $2.2 \times 10^{-16}$ | Small constant added to $\|v\|$ in the denominator of Eq. (26) to avoid division by zero. |
| Maximum Step Length | $s_{max}$ | $1.2 m$ | Upper bound on forward foot displacement per step. |

| | | | |
|---|---|---|---|
| Vertical CoM Oscillation Amplitude | $z_{amp}$ | $0.03m$ | Amplitude of sinusoidal vertical CoM bob per gait cycle. |

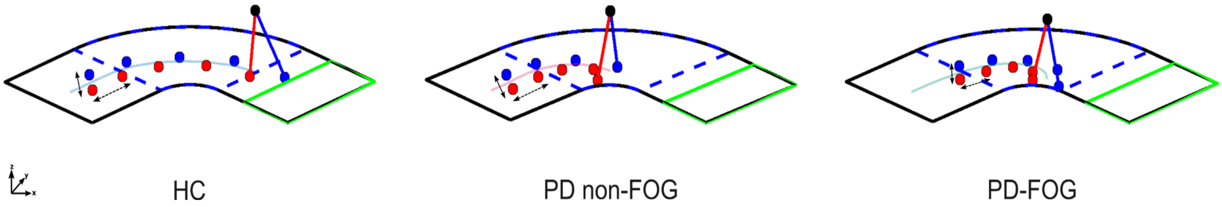


**Supplementary Figure S1. Turning strategies across conditions:** Simulated real-time foot placements during a 90° turn for HC, PD non-FOG, and PD-FOG. The black marker denotes the center of mass (CoM), while blue and red markers indicate the left and right foot placements, respectively. The solid and dashed double-headed arrows represent step width and step length, respectively and the agent's path is drawn over the track for all cases. The visualization captures the agent's current posture alongside a history of ~10 previous steps generated by the gait model. Comparing the trajectories across the three conditions shows a clear progressive reduction in step length from HC to PD-FOG along with a moderate reduction in step width between HC and the PD groups.

### Task Learning and Overall Success Rates

To evaluate the learning capacity of the agents, we tracked the overall success rate — defined as successfully reaching the final reward box, across 50 discrete evaluation blocks (10 trials per block). Supplementary Fig. S2 illustrates these learning curves for each turn angle, highlighting the progression from initial exploration (unshaded region) to the final low-noise testing phase (green shaded region). Healthy Controls demonstrated a robust ability to learn the turn task. Across most turn angles, the HC success rate climbed steadily during the training phase as the agent optimized its value function. In the testing phase where the highly variable, gradient-driven exploration is reduced to a minimal baseline noise, the HC agent's performance reliably peaked, frequently exceeding its training-phase success rates. While the extreme 180° turn proved more difficult, the HC agent still displayed a clear, upward learning trajectory.

Conversely, simulating dopamine depletion severely affected the learning architecture in both PD models. For both the PD non-FOG and PD-FOG agents, the learning curves remained almost completely flat throughout the entire simulation. Except for the 0° turn, the success rate for the PD agents hovered around 1 to 3 successful trials per block, with no meaningful improvement transitioning into the testing phase.

This striking lack of progression confirms that clamping the TD-error fundamentally prevents the agent from accumulating positive experiences and updating its policy. Unable to properly learn the value gradient of the track, the PD agents are left to rely on impaired, trial-and-error navigation, explaining the lower success rates and severe trajectory deviations observed in the behavioral data as seen in Fig. 3a.

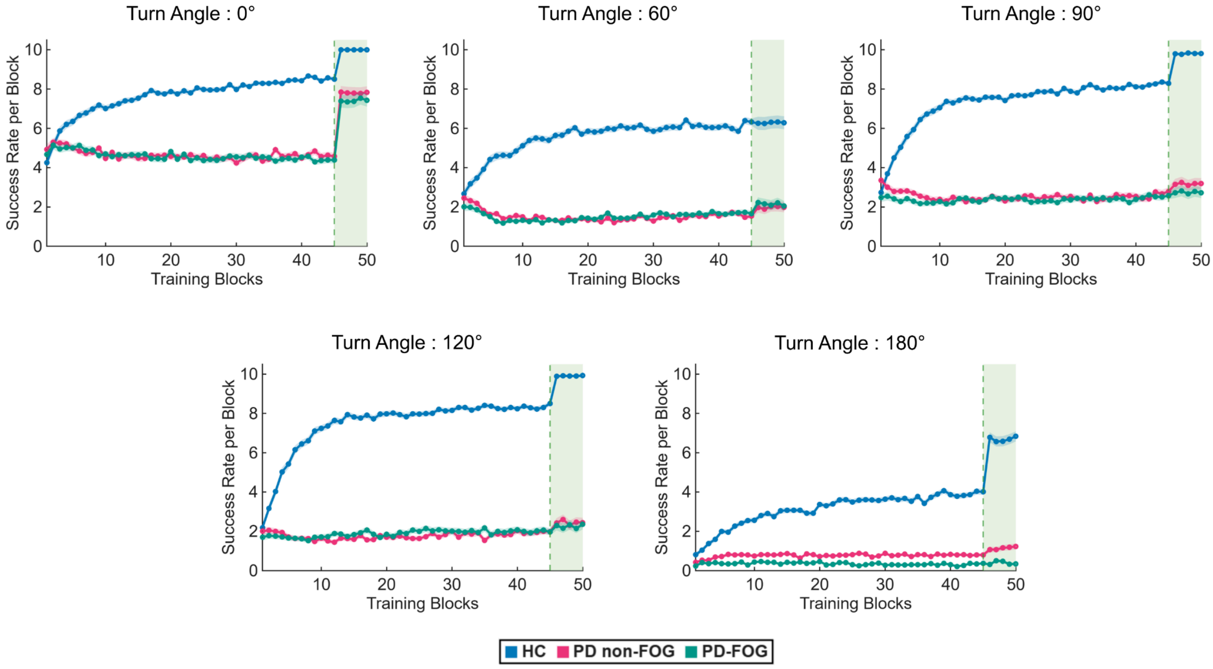

**Supplementary Figure S2. Task Success Rates Across Turn Angles:** Success rate (out of 10 trials per block), plotted across 50 training blocks for HC, PD non-FOG, and PD-FOG at each turn angle. Solid lines denote group means, while shaded error bands represent $\pm$ 1 standard error of the mean (SEM). The unshaded region indicates the training phase (stochastic policy, exploration active), while the green-shaded region (blocks 46–50) indicates the low-noise testing phase (minimal exploration noise). HC success rate increases steadily during training, however both PD groups plateau at a substantially lower success rate throughout training. Neither PD group approaches HC's performance during the testing phase at any given turn angle.

**Parameter Space Analysis: Velocity Heatmaps**

To investigate the interaction between dopaminergic depletion and exploration, the agent's velocity profiles were evaluated across a 2D grid mapping the TD-error clamp limit $\delta_{lim}$ against the exploration amplitude $Ae_{eff}$. The velocity heatmaps show the agent's mean turning speed across the turn during testing. As with value in Fig. 2c, $\delta_{lim}$ was the primary determinant of velocity. From 60° to 120°, velocity increased sharply between $\delta_{lim}$= 0.001 and 0.01, then more gradually, reaching its maximum at $\delta_{lim}$ = 1. At 180°, velocity peaked at $\delta_{lim}$ = 0.1, while at 0°, it was highest at $\delta_{lim}$ = 1. The very low velocities at small $\delta_{lim}$ are consistent with $\delta_{lim}$ - dependent scaling of the Go/No-go action gains in the PD condition, which suppresses movement. Since the gain scaling uses $Angle_{ratio}$, which is 0 at 0°, this suppression vanishes there. The influence of $Ae_{eff}$ was weaker. At 0° and $\delta_{lim}$= 1, higher $Ae_{eff}$ strongly reduced velocity, consistent with excessive exploration during learning being detrimental for simple paths. From 60° to 120°, $Ae_{eff}$ produced only small changes. At 180°, the highest velocities were seen for $Ae_{eff}$= 1.5–2.5 and $\delta_{lim}$= 0.1, suggesting that higher exploration may be beneficial for the sharpest turn.

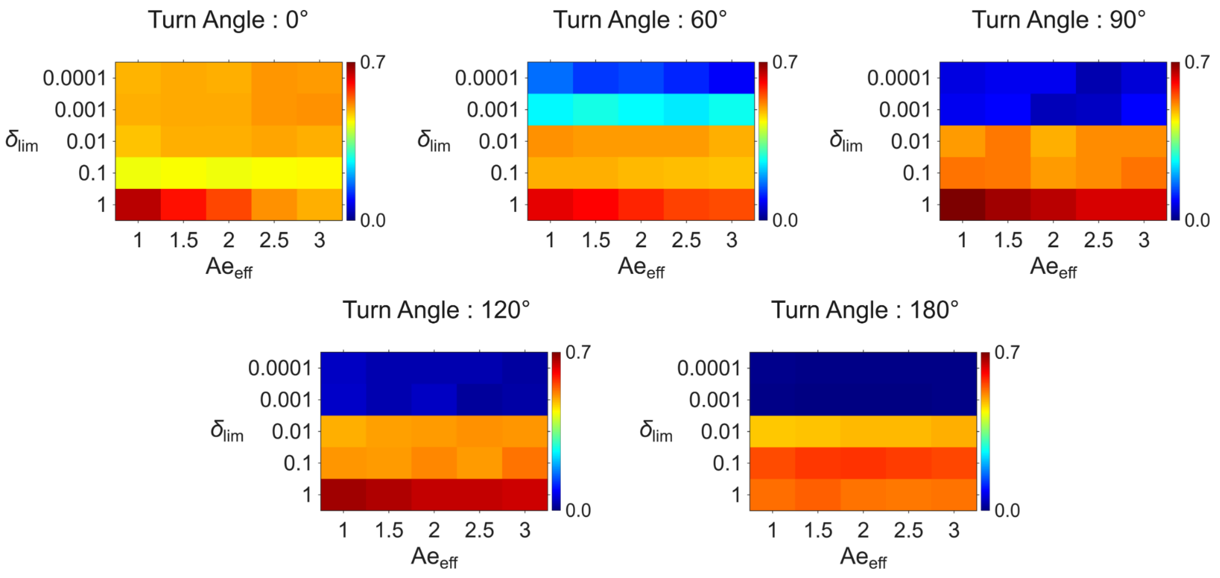


**Supplementary Figure S3. Parameter Space Heatmaps for Velocity:** Mean turning velocity heatmaps along the agent's test trajectories, mapped across the TD-error clamp limit ($\delta_{lim}$; y-axis, 0.0001–1) and the exploration amplitude ($Ae_{eff}$; x-axis, 1–3), shown for 0°, 60°, 90°, 120°, and 180° turn angles. Warmer colors indicate higher velocities; color scales are directly comparable across angles. Velocity is strongly modulated by $\delta_{lim}$, being lowest for very low $\delta_{lim}$ for most angles, whereas $Ae_{eff}$ has a smaller effect.
